# Inhibitors of trehalose-6-phosphate synthase activity in fungal pathogens compromise thermal tolerance pathways

**DOI:** 10.1101/2025.03.07.642065

**Authors:** Yi Miao, Vikas Yadav, William Shadrick, Jiuyu Liu, Alexander R. Jenner, Connie B. Nichols, Clifford Gee, Martin Schäfer, Jennifer L. Tenor, John R. Perfect, Richard E. Lee, Richard G. Brennan, Erica J. Washington

**Author notes:** Present address: Division of Life Science, The Hong Kong University of Science and Technology, Clear Water Bay, Kowloon, Hong Kong SAR, China. Present address: Incyte Corporation, Wilmington, Delaware (William Shadrick). Present address: Department of Chemistry and Biochemistry, Creighton University, Omaha, Nebraska, USA. Present address: GCP-Service International Ltd & Co. KG, Bremen, Germany. Co-first authors (Yi Miao, Vikas Yadav).

## Abstract

Infections caused by fungal pathogens such as *Candida* and *Cryptococcus* are associated with high mortality rates, partly due to limitations in the current antifungal arsenal. This highlights the need for antifungal drug targets with novel mechanisms of action. The trehalose biosynthesis pathway is a promising antifungal drug target because trehalose biosynthesis is essential for virulence in *Cryptococcus neoformans* and *Candida albicans* and is also a mediator of fungal stress responses, such as thermotolerance. To exploit its untapped antifungal potentials, we screened the St. Jude 3-point pharmacophore library to identify small molecule inhibitors of the first enzyme in the trehalose biosynthesis pathway, trehalose-6-phosphate synthase (Tps1). Structure-guided optimization of a potent hit, SJ6675, yielded a water-soluble inhibitor named 4456dh. Employing biochemical, structural and cell-based assays, we demonstrate that 4456dh inhibits Tps1 enzymatic activity, suppresses trehalose synthesis and exerts a fungicidal effect. Notably, the structure of Tps1 in complex with 4456 reveals that 4456 occupies the substrate binding pocket. Importantly, 4456dh renders normally thermotolerant fungal pathogens unable to survive at elevated temperatures, which is critical as we investigate the emergence of fungi from the environment due to a warming climate. Overall, this work develops the water-soluble 4456dh as an early-stage antifungal drug that has a distinct mechanism of action compared to existing clinical antifungals.

**Importance:** The rise of fungal infections in recent years is alarming due to an increase in the vulnerable immunocompromised population, global temperature increase and limited antifungal treatment options. One of the major hurdles in developing new drugs is the identification of fungal-specific antifungal drug targets due to highly conserved cellular machinery between fungi and humans. Here, we describe a small molecule inhibitor, 4456dh, of the trehalose biosynthesis pathway. This pathway is present in fungi but not humans. Trehalose plays a critical role in stress responses such as thermotolerance in fungal pathogens and is essential for their virulence. We show that treatment with 4456dh blocks production of trehalose and renders fungal cells inviable. Thus far, 4456dh is active against two fungal pathogens of critical importance suggesting a broad-spectrum activity.

## Introduction

The contribution of fungal pathogens to human mortality goes greatly unnoticed compared to that of viruses and bacteria and even other eukaryotic pathogens. Remarkably, the combined mortality associated with fungal pathogens is similar to that caused by malaria and tuberculosis (1). Each year a diverse collection of fungal pathogens cause diseases that result in approximately 6.5 million infections and 2.5 million directly attributable deaths worldwide (2). The World Health Organization recently documented the significance of these fungal pathogens and listed *Cryptococcus neoformans* and *Candida albicans* as critical pathogens, amongst others, and made the call for increased development of antifungal therapeutics, fungal surveillance and diagnostics (3).

The current antifungal drug armamentarium consists of polyenes, azoles and echinocandins. While these antifungal drugs have proven to be powerful, they also have drawbacks that include the lack of bioavailability, lack of effectiveness against multiple pathogens and an increase in the likelihood of antifungal drug resistance (4). Current antifungal drugs are also often toxic and require long and difficult treatment regimens (5). Therefore, these drugs are insufficient to control the current state of fungal infections. Moreover, new fungal pathogens, such as the multidrug-resistant thermotolerant yeast *Candida auris*, are emerging due to a warming climate (6–11).

Because fungi and humans are both eukaryotes, it is difficult yet advantageous to identify an antifungal drug target that is not conserved between both species. An antifungal drug with a target not found in humans is likely to have fewer off-target effects and be less toxic for the immunocompromised patients who are susceptible to invasive fungal infections. Other features of antifungal drugs that are desirable include the possibility of being broad-spectrum in the case of poor diagnostics and a low likelihood that resistance to the antifungal drug will develop.

The trehalose biosynthesis pathway in fungal pathogens has been described as a promising antifungal drug target (12). The trehalose biosynthesis pathway is conserved amongst insects, plants, bacteria and fungi (13). However, the trehalose biosynthesis pathway is not found in mammals (12, 13). Given that the key components of the trehalose biosynthesis pathway are structurally conserved, it may be possible to generate a broad-spectrum antifungal drug that targets its components. Attempts have been made to generate compounds that inhibit trehalose biosynthesis in insects (14). Closantel was identified as a potential bioactive molecule that targets the trehalose biosynthesis pathway and has potent activity against *C. neoformans* and *Cryptococcus gattii in vitro*. However, closantel does not have antifungal activity in murine models (12).

The trehalose biosynthesis pathway in fungal pathogens is a two-step process initiated when trehalose-6-phosphate synthase, Tps1, converts uridine-diphosphate-glucose (UDPG) and glucose-6-phosphate (G6P) to trehalose-6-phosphate (T6P) (15). T6P is dephosphorylated by the trehalose-6-phosphate phosphatase (Tps2) to produce trehalose (15). Several pathogens also contain additional trehalose biosynthesis proteins. For example, in *Aspergillus fumigatus* there are apparently multiple Tps1 enzymes (16). In ascomycetes, including *A. fumigatus*, *C. albicans* and *S. cerevisiae,* there are regulatory trehalose biosynthesis proteins, which lack catalytic activity and are thought to act as scaffolds to enable complex formation of the trehalose biosynthesis proteins (17).

Trehalose is a nonreducing disaccharide composed of two glucose molecules linked by an α,α-1,1-glycosidic bond (α-D-glucopyranosyl-(1→1)-α-D-glucopyranoside). Trehalose is a highly stable molecule that can help proteins and cellular membranes withstand dehydration, salinity, and heat (18–20). Trehalose can also serve as an energy source and is broken down by trehalase into two glucose molecules that funnel into energy-generating pathways such as glycolysis and the pentose phosphate pathway (15, 21, 22). Importantly, the trehalose biosynthesis pathway is strongly associated with virulence in *C. neoformans* and *C. albicans*. In the absence of *TPS1* and *TPS2*, *Cryptococcus* and *Candida* species cannot survive in mammalian hosts (23–27). The contribution of trehalose biosynthesis to thermotolerance in *C. albicans* and *C*. *neoformans* is critical. The inhibition of trehalose biosynthesis may help control the expansion of thermotolerant environmental isolates, which have the potential to become pathogens.

We now have the tools to take a structure-based approach to developing an inhibitor of Tps1. The recent determination of structures of trehalose biosynthesis proteins from fungal pathogens, including *C*. *albicans* and *A*. *fumigatus* has enabled us to utilize a structure-guided approach to designing inhibitors of trehalose biosynthesis proteins (28, 29). Most recently, cryo-electron microscopy was used to determine detailed binding residues in the *C. neoformans* substrate-binding pocket (30). Tps1 is a GT-B family, retaining glycosyltransferase and contains two catalytic subdomains, connected by a kinked, C-terminal α-helix (31). The N-terminal subdomain binds the sugar acceptor, G6P, and the C-terminal subdomain contains the donor, UDPG. Given that Tps1 utilizes a substrate-assisted mechanism of catalysis, proper alignment of each substrate is required for the formation of T6P. Therefore, targeting the substrate-binding pocket is a critical and facile route to disrupt the function of Tps1. Additionally, the ability to generate large amounts of pure recombinant *C. neoformans* Tps1 and *C. albicans* Tps1 expressed and purified in *E. coli* has made high-throughput compound screens possible.

In this work, we developed a fluorescence polarization-based high-throughput screening platform and screened a chemical library containing 750 compounds to identify Tps1 inhibitors. We determined the structure of a top hit bound to *C. albicans* Tps1. Guided by this structural information, we rationally optimized the hit and developed a water-soluble derivative, 4456dh. 4456dh demonstrates broad-spectrum inhibitory effects against multiple pathogenic fungal species, including *Candida* and *Cryptococcus*. Furthermore, 4456dh displays temperature-dependent fungicidal activity, phenocopying the *tps1*Δ fungicidal phenotype. Finally, we show that in fungal cells 4456dh inhibits the biosynthesis of trehalose. These data indicate that the mechanism of action of 4456dh is indeed trehalose-dependent and fully exploits the “druggable” thermotolerant characteristics of various pathogenic fungal species.

Overall, this work highlights the combination of high-throughput screening, structure-guided hit optimization and cellular assays to develop a promising water-soluble small molecule with desired antifungal properties and the potential for development of a new antifungal therapeutic.

## Results

### Seven compounds were identified from a high-throughput screen to inhibit CaTps1 activity

As the first step of the trehalose biosynthesis pathway, Tps1 synthesizes T6P from the substrates UDP-glucose and G6P followed by the release of UDP and T6P (15). To identify a small molecule inhibitor of Tps1 activity, we adapted a high-throughput screen (HTS) utilizing the UDP Transcreener Fluorescence Polarization (FP) assay (32, 33). The FP assay detects the UDP released as a result of the Tps1-mediated enzymatic reaction (Fig. 1A). The FP assay Z’ factor, which describes the suitability of an assay for a HTS, is 0.87, indicating an ideal signal-to-noise ratio (Fig. S1A) (34). There was a greater than 175 millipolarization (mP) shift between the positive control novobiocin, an inhibitor with activity against a bacterial trehalose-P-synthase, and the negative control DMSO (Fig. S1A) (35). Recombinant Tps1 from *C. albicans*, hereafter referred to as CaTps1, was expressed in *E. coli* and purified via an N-terminal 6xHis-tag and Ni^2+-^NTA affinity chromatography, followed by size exclusion chromatography (Fig. 1B). A 3-point pharmacophore (3PP) library, which consisted of a subset of 750 compounds of less than 300 Da in molecular mass from the St. Jude Children’s Research Hospital chemical library, was screened for the ability to inhibit 6xHis-CaTps1 activity (36). In the primary screen, 32 (4.3 %) compounds that inhibited the activity of 6xHis-CaTps1 by more than 10% at 500 µM concentration were identified (Fig. 1C,D). A 10% cutoff provided significant number of hits to validate using a dose response assay (Fig. S1B). At the completion of the screen, seven compounds that inhibit the activity of CaTps1 in a dose-dependent manner were selected for further investigation (Fig. 1E and Table 1).

**Figure 1.**
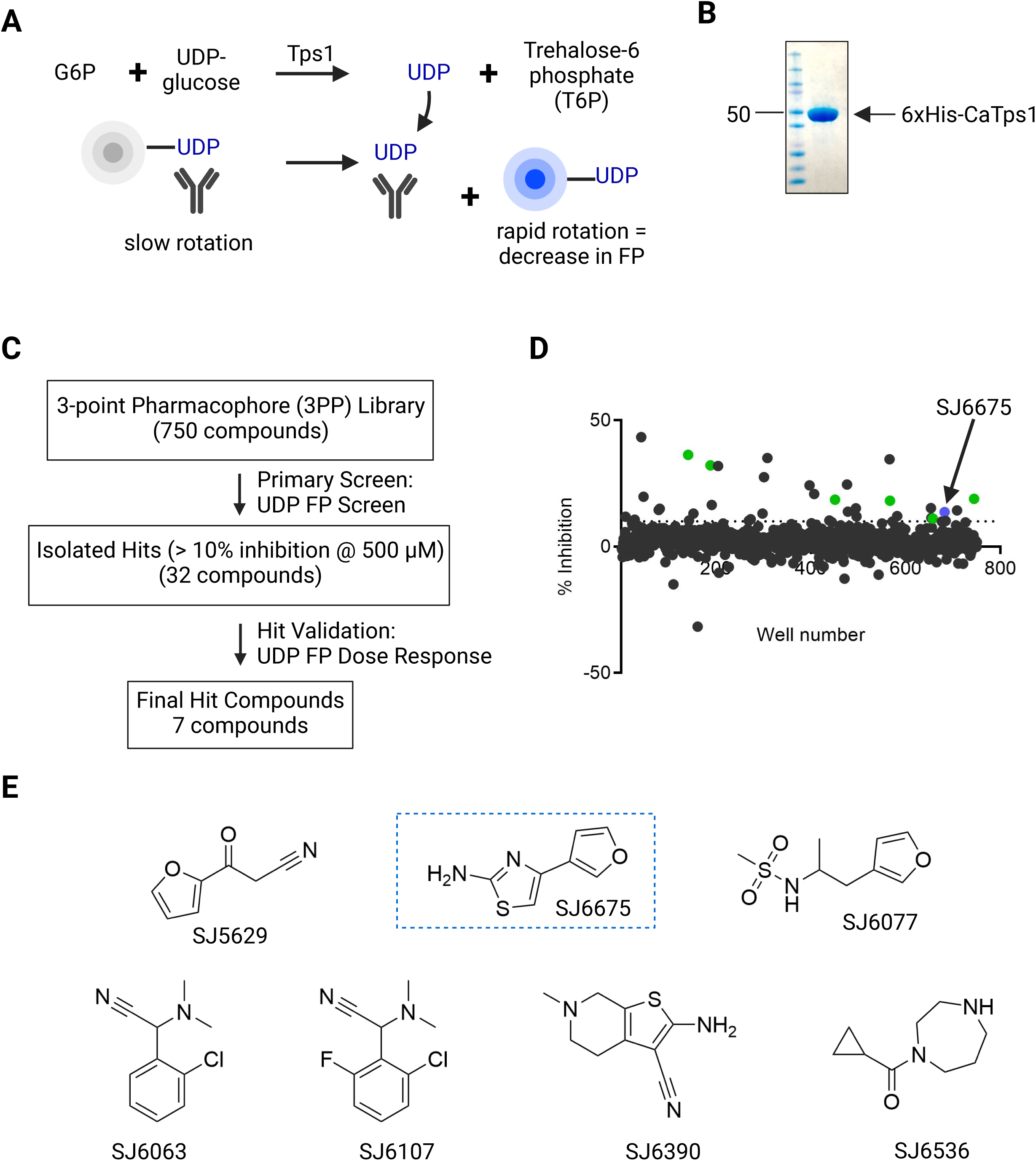
Seven Tps1 inhibitors were identified using a high-throughput fluorescence polarization screen. **A)** Diagram of the Transcreener assay for Tps1 activity. **B)** Coomassie-stained SDS polyacrylamide gel of purified recombinant 6xHis-CaTps1. Molecular weight markers are shown with the 50 kDa protein standard highlighted with an arrow. **C)** Pipeline and results from the Tps1 UDP Transcreener assay. **D)** Dot plot of % inhibition and compounds that were tested for inhibition of 6xHis-CaTps1 activity. The 32 compounds above 10% inhibition at 500 µM are highlighted. Highlighted in green are 6 of the compounds that resulted in dose-dependent inhibition of 10% or greater of 6xHis-CaTps1 activity. In blue and highlighted with an arrow is SJ6675. **E)** The chemical structures and identities of validated 6xHis-CaTps1-inhibiting hit compounds. The compound SJ6675 used in additional studies is boxed.

**Table 1.** Small molecules identified in high-throughput screen that inhibit the activity of 6xHis-CaTps1.

| Compound | Chemical Formula | Molecular Weight | CaTps1 FP (% inhibition) | CaTps1 Inhibition Assay (μM) |
| --- | --- | --- | --- | --- |
| SJ000805629-1 | C<sub>7</sub>H<sub>5</sub>NO<sub>2</sub> | 135.12 | 32.19 | 685.4 |
| SJ000866675-1 | C<sub>7</sub>H<sub>6</sub>N<sub>2</sub>OS | 166.20 | 13.69 | 983.3 |
| SJ000866077-1 | C<sub>8</sub>H<sub>13</sub>NO<sub>3</sub>S | 203.26 | 11.12 | 1231.5 |
| SJ000866063-1 | C<sub>10</sub>H<sub>11</sub>CIN<sub>2</sub> | 194.66 | 18.96 | 767.5 |
| SJ000866107-1 | C<sub>11</sub>H<sub>13</sub>CIN<sub>2</sub> | 208.69 | 18.24 | 1152.3 |
| SJ000866390-1 | C<sub>10</sub>H<sub>13</sub>N<sub>3</sub> | 175.24 | 36.36 | 608.7 |
| SJ000866536-1 | C<sub>9</sub>H<sub>16</sub>N<sub>2</sub>O | 168.24 | 18.64 | 546.9 |

### Generation of a library of derivatives of SJ6675

We initiated a structure-guided approach to generate derivatives of the compounds by attempting *de novo* co-crystallization of each of the seven compounds with CaTps1. We successfully obtained a 3.5 Å resolution structure of SJ6675 ((4-furan-3-yl)thiazol-2-amine) in complex with CaTps1 (Fig. S2A and Table S1). This structure revealed the presence of SJ6675 in the substrate-binding pocket of CaTps1 (Fig. S2A), proximal to conserved substrate-binding residues R280, K285, N382 and L383 (28). These data indicate that SJ6675 likely interferes competitively with the binding of the UDPG substrate.

We designed a library of SJ6675 derivatives and tested the derivatives for inhibition of CaTps1 activity (Fig. S2B,C). Many of the compounds, diluted in DMSO, precipitated when added to the Tps1 activity assay reaction and several of the compounds increased the activity of CaTps1 (Fig. S2C). However, one of these compounds, 4456, significantly inhibited the activity of CaTps1 and did not precipitate in the assay solution (Fig. S2C).

### 4456dh binds Tps1 with high affinity and inhibits the Tps1-mediated release of UDP

To determine the ability to use 4456 as a lead compound using biochemical approaches, we generated a water-soluble derivative of 4456, which we shall now refer to as 4456dh, for 4456-dichloride hydrate (Fig. 2A). The composition of 4456dh was confirmed by NMR and elemental analysis (Fig. S3).

**Figure 2.**
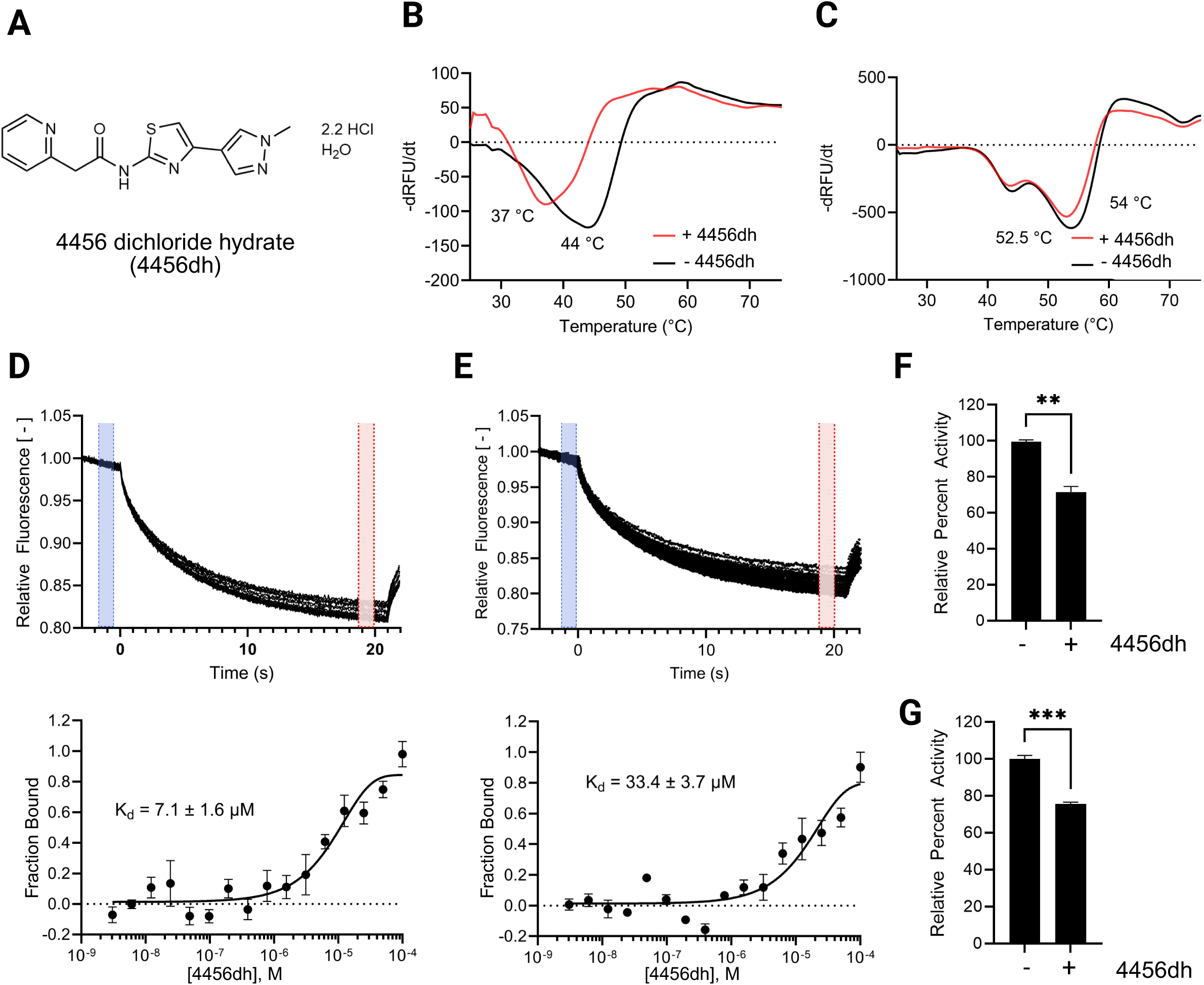
4456dh binds Tps1 and inhibits Tps1 activity. **A)** Structure of 4456dh. **B-C)** Glo-melt thermal shift assays reveal 4456dh destabilizes 6xHis-CaTps1 (B) and 6xHis-CnTps1 (C). The first derivative of the raw data was used to determine the melting temperature of 1 µM 6xHis-CaTps1 (B) and 1 µM 6xHis-CnTps1 (C) treated without (black) or with 2 mM 4456dh (red). **D-E)** Microscale thermophoresis assays demonstrate 4456dh binds to 6xHis-CaTps1 (D) and 6xHis-CnTps1 (E) with indicated K_d_. The top panel shows a representative of the MST trace with the cold region labeled highlighted in blue and the hot region highlighted in red. The bottom panel shows the dose-response curve. The resulting dose-response curves were fitted to a one-site binding model to extract K_d_ values; the standard deviation was calculated using the K_d_ values from each independent experiment (n=3). **F)** Tps1 enzyme activity assay with 5 µM 6xHis-CaTps1 (F) and 6xHis-CnTps1 (G) +/- 1 mM 4456dh. Error bars represent standard error. Comparison to control, untreated 6xHis-CaTps1, was performed using Student’s t-test; **p < 0.01, *** p< 0.001.

Thermal shift assays were completed to determine if 4456dh binds Tps1. We hypothesized that the binding of 4456dh to Tps1 would increase Tps1 stability, resulting in a higher melting temperature. Therefore, CaTps1 and *C. neoformans* Tps1, hereafter referred to as CnTps1, were incubated with 4456dh overnight at 4 °C. Glo-Melt dye was added to the mixture prior to determining the thermostability. Surprisingly, we observed that 4456dh decreased the melting temperature of both CaTps1 and CnTps1 in thermal shift assays indicating a destabilization of the Tps1 enzyme upon binding (Fig. 2B,C). These data indicate that 4456dh binds to both CaTps1 and CnTps1 but leads to destabilization of the Tps1 enzyme.

To determine the binding affinity of 4456dh and CaTps1 and CnTps1, microscale thermophoresis (MST) was used. Given the intrinsic fluorescence of CaTps1 and CnTps1, we were able to detect binding without the use of additional fluorophores. MST experiments revealed that the binding affinities of 4456dh with recombinant 6xHis-CaTps1 and 6xHis-CnTps1 are 7.1 µM and 33.4 µM, respectively (Fig. 2D,E). These binding affinities are consistent with the concentration of 4456dh at which we can detect a shift in Tps1 stability in the thermal shift assays. The ability of 4456dh to inhibit the activity of CaTps1 was tested using a spectrophotometric-coupled Tps1 enzyme assay, in which recombinant 6xHis-CaTps1 was incubated with 1 mM 4456dh at 4 °C for 16 hours. The activity of 6xHis-CaTps1 was reduced approximately 30% in the presence of 4456dh (Fig. 2F). Similarly, the effect of 1 mM 4456dh on 6xHis-CnTps1 activity was determined. 4456dh inhibited the activity of 6xHis-CnTps1 activity by approximately 30% (Fig. 2G). Therefore, 4456dh directly binds and inhibits the activity of Tps1 from *C. albicans* and *C. neoformans*.

### 4456dh binds the substrate-binding pocket of CaTps1

To gain insight into the mode of action of 4456dh, we grew crystals of 6xHis-tagged *C. albicans* Tps1 (6xHis-CaTps1) in complex with 4456 and determined the structure to a moderate 3.35 Å resolution (Fig. 3A and Table 2). The structure of the 6xHis-CaTps1-4456 complex was solved using molecular replacement and the structure of CaTps1 bound to UDP and G6P (PDB: 5HUU), with UDP and G6P removed as the search query. The final model of the 6xHis-CaTps1-4456 complex consists of N-terminal and C-terminal subdomains comprising residues 6 to 239 and 246 to 437, respectively, followed by the C-terminal α-helix (Fig. 3A). The final model also contains one 4456 molecule per subunit of 6xHis-CaTps1 (Fig. 3A,B).

**Figure 3.**
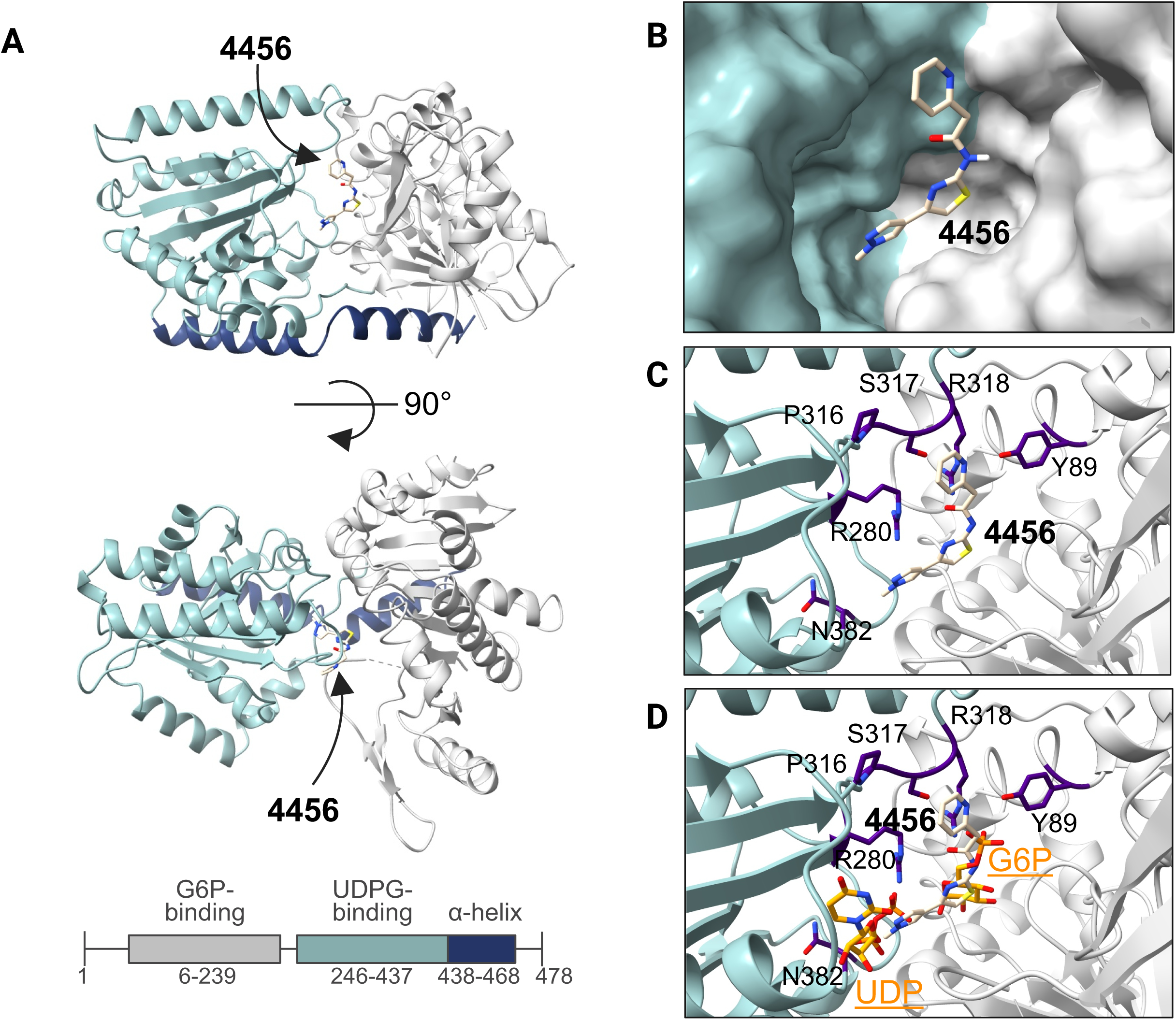
Structure of *C. albicans* Tps1 in complex with 4456. **A)** One subunit of the CaTps1-4456 complex shown as a ribbon diagram. **B)** Surface-rendered structure revealing 4456, shown as atomic-colored sticks, binds in the deep cavity of the substrate-binding pocket proximal to known substrate-binding residues. **C)** Tps1 residues within 4 Å of 4456 are labelled, shown as stick representations and indigo colored. **D)** Residues within 4 Å of 4456 are proximal to residues that also interact with product and substrate, UDP and G6P (atom-colored sticks), in the CaTps1-UDP-G6P complex (PDB: 5HUU) (28).

**Table 2.** Data collection and refinement statistics of structure of 6xHis-CaTps1 in complex with 4456.

| **6xHisCaTps1 – 4456 complex** |  |
| --- | --- |
| Resolution range | 58.78 - 3.35 (3.47 - 3.35) |
| Space group | P 6<sub>4</sub> 2 2 |
| Unit cell | 117.6 117.6 134.0 90 90 120 |
| Total reflections | 75,865 (7,778) |
| Unique reflections | 7,644 (753) |
| Multiplicity | 9.9 (10.3) |
| Completeness (%) | 91.66 (94.0) |
| Mean I/sigma(I) | 14.9 (3.3) |
| Wilson B-factor | 78.7 |
| R-merge | 0.1037 (0.2833) |
| R-meas | 0.1089 (0.297) |
| R-pim | 0.03209 (0.08654) |
| CC<sub>1/2</sub> | 0.998 (0.932) |
| CC* | 0.999 (0.982) |
| Reflections used in refinement | 7641 (753) |
| Reflections used for R-free | 716 (71) |
| R-work | 0.2504 (0.3147) |
| R-free | 0.2893 (0.3509) |
| Number of non-hydrogen atoms | 3644 |
| macromolecules | 3623 |
| ligands | 21 |
| solvent | 0 |
| Protein residues | 456 |
| RMS(bonds) | 0.003 |
| RMS(angles) | 0.53 |
| Ramachandran favored (%) | 94.9 |
| Ramachandran allowed (%) | 4.45 |
| Ramachandran outliers (%) | 0.67 |
| Rotamer outliers (%) | 0.00 |
| Clashscore | 5.12 |
| Average B-factor | 83.35 |
| macromolecules | 83.25 |
| ligands | 100.62 |

Statistics for the highest-resolution shell are shown in parentheses.

During refinement, electron density corresponding to 4456 was identified in the substrate-binding pocket of CaTps1 between the N-terminal G6P-binding lobe and the C-terminal UDPG-binding lobe. Polder omit maps, which exclude bulk solvent to allow for detection of weak density, of the bound inhibitor reveal the binding of 4456 (Fig. S4) (37). 4456 was modelled into the electron density between these two lobes and was found to make contacts with multiple conserved residues and functionally critical substrate-binding residues (Fig. 3C and Fig. S5) (28). The conserved residues near 4456 (Y89, R280, P316, S317, R318 and N382) are proximal to the native product, UDP, and substrate G6P (Fig. 3D) that are required for catalytic activity. For example, the hydroxyl group of the side chain of Y89 makes hydrophilic interactions with the G6P phosphate moiety and mutation of Y89 to phenylalanine results the elimination of Tps1 activity (28). The model of the catalytic modified-Rossman fold domains and the C-terminal α-helix is structurally similar to previously determined structures of Tps1 from *C. neoformans*, *A. fumigatus* and *E. coli,* and therefore, demonstrate the likelihood that 4456 also binds in the conserved catalytic pockets of Tps1 from these additional organisms (Fig. S6A) (28, 30). Using a ConSurf analysis, the degree of conservation was mapped onto Tps1 structures and revealed that the most highly conserved residues are either buried in the substrate-binding pocket or are proximal to that region (Fig. S6B) (38). This finding was expected given that 4456 is a derivative of a lead compound that also binds the substrate-binding pocket of CaTps1 (Fig. S2).

### 4456dh is bioactive against Candida and Cryptococcus cells

We tested the effects of 4456dh on live cells by determining the minimal inhibitory concentration causing 80% growth reduction (MIC_80_). Wild-type *C. albicans* and *C. auris* cells were incubated with 4456dh in yeast-peptone-dextrose (YPD) medium at 42 °C and 39 °C, respectively (Fig. 4A). 4456dh inhibited the growth of *C. albicans* at 42 °C with an MIC_80_ of 13 mM, whereas no MIC was detected at 30 °C (Figure 4A and Fig. S7A). 4456dh inhibited the growth of *C. auris* with an MIC_80_ of 7 mM at 39 °C (Fig. 4A). The MIC_80_ increased at 30 °C to greater than 13 mM (Fig. S7A). We characterized the fungicidal activity of 4456dh by growing the *C. albicans* on compound-free YPD plates after exposure to 4456dh and observed temperature-dependent fungicidal activity of 4456dh on *C. albicans* at 42 °C (Fig. 4A and Fig. S7B). Interestingly, fungicidal activity of 4456dh on *C. auris* was also detected at both 30 °C and 39 °C (Fig. 4A and Fig. S7B).

**Figure 4.**
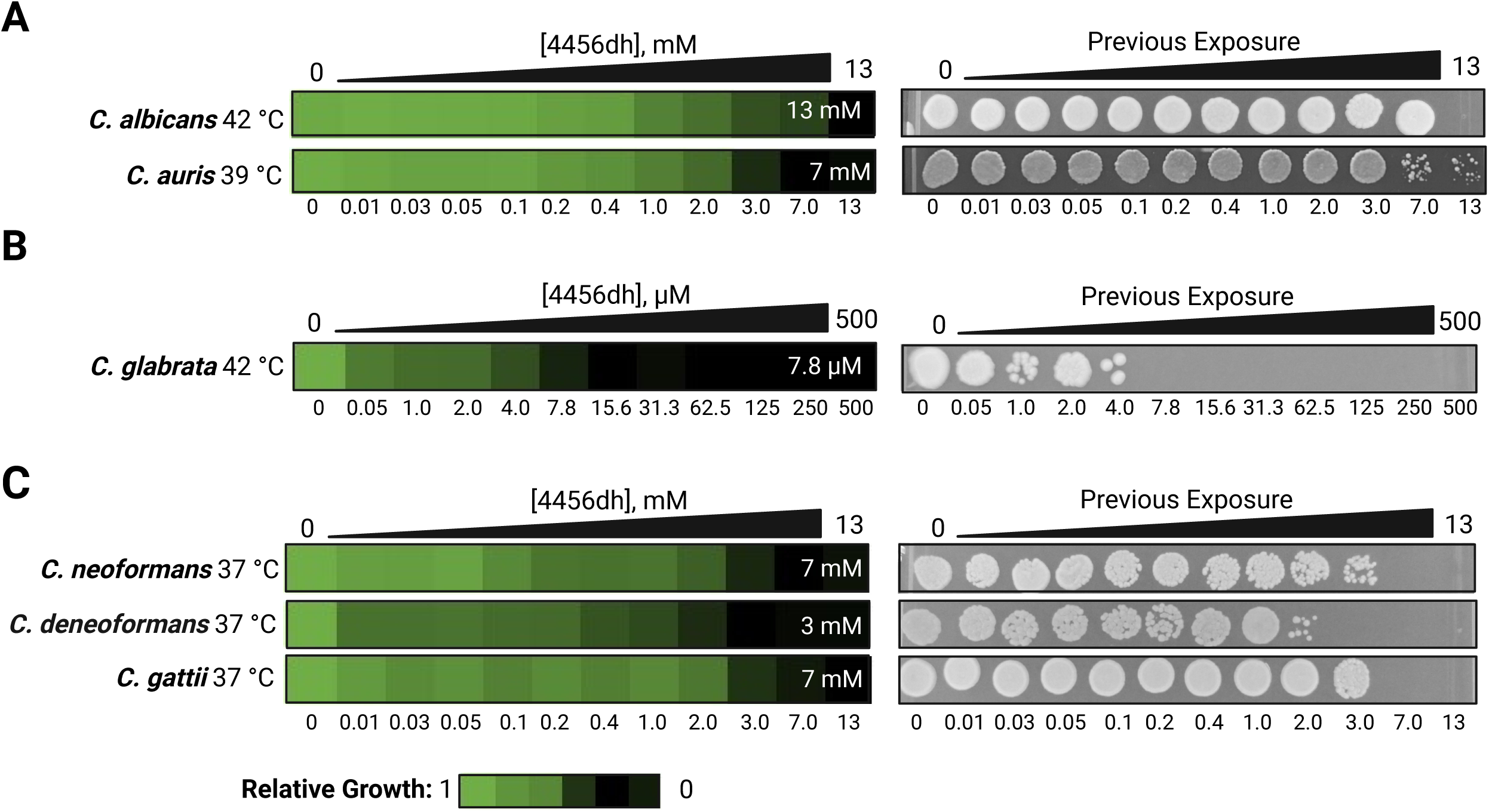
4456dh inhibits the growth of *Candida* and *Cryptococcus* species. Two-fold dose response assays were performed against **A)** *C. albicans* at 42 °C and *C. auris* at 39 °C, **B)** *C. glabrata* at 42 °C, and **C)** *C. neoformans*, *C. deneoformans* and *C. gattii* at 37 °C. 4456dh was administered at the concentrations indicated. Cultures were grown in YPD at the listed temperatures for 48 hours for *Candida* species and 72 hours for *Cryptococcus* species. Experimental temperatures reflect the temperatures at which the fungi experience heat stress. (Left) Relative growth of treated samples was calculated by averaging technical triplicates and normalizing the OD_600_ of each treated well to the average OD_600_ of the untreated wild-type controls, after the absorbance due to the compound was subtracted. The MIC_80_ values, calculated based on the OD_600_ values, are indicated in the white text on each heat map. (Right) Representative spot assays show the ability of cells to grow after exposure to 4456dh and indicate fungicidal activity of 4456dh.

4456dh displayed the most potent activity on *C. glabrata*, with an MIC_80_ of 7.8 µM at 42°C (Fig. 4B). Furthermore, there was significant fungicidal activity of 4456dh on *C. glabrata* in concentrations starting at approximately 7.8 µM (Fig. 4B). The activity against *C. glabrata* was specific to 42 °C, as no MIC or fungicidal activity was detected at 30 °C (Fig. S7A,B).

We next tested the activity of 4456dh on *Cryptococcus* species at 37 °C. We found that 4456dh inhibited the growth of *C. neoformans*, *C*. *deneoformans*, and *C. gattii* with MIC_80_ of 7 mM, 3 mM, and 7 mM, respectively (Fig. 4C). Fungicidal activity of 4456dh was also detected in all the *Cryptococcus* species tested at 37 °C (Fig. 4C). Interestingly, 4456dh also displayed fungicidal activity on *C. deneoformans* at 30 °C at 13 mM (Fig. S7C).

Given that *C. neoformans tps1*Δ yeasts are fungicidal at elevated temperature and not at 30 °C, the fungicidal activity of 4456dh at 30 °C indicated that in addition to Tps1 there is another target of 4456dh. To test this hypothesis, we performed zone of inhibition studies in the *C. deneoformans* background (Fig. S8A). At 37 °C, there was a distinct zone of inhibition surrounding 4456dh (Fig. S8A). Consistent with our prediction of an additional target of 4456dh, we detected a diffuse, yet visible zone of inhibition due to 4456dh on *C. deneoformans tps1*Δ cells (Fig. S8B). Inhibitors of Tps1 may work synergistically with known antifungal drugs. In *C. glabrata* the trehalose biosynthesis mutant, *tps2*Δ, is more susceptible to fluconazole at 37 °C (39). Therefore, we hypothesized that there might be synergy between 4456dh and fluconazole. We performed Epsilometer tests (E-tests) and detected synergy between 4456dh and fluconazole in *C. neoformans H99* in a dose-dependent manner (Fig. S9).

### 4456dh inhibition mimics loss of functional Tps1

Given that 4456dh inhibits the activity of CaTps1 and CnTps1 *in vitro* and has bioactivity against multiple *Candida* and *Cryptococcus* species, we tested if 4456dh affects trehalose production in cells. We quantified trehalose levels in cells treated with 4456dh at sub-lethal concentrations as determined by the MIC assays using a colorimetric spectrophotometric assay. We measured trehalose production in wild-type *C. albicans* at 30 °C and 42 °C, in the presence and absence of 4456dh. We observed that at 30 °C, there was a minimal amount of trehalose detected and, therefore, there was no significant change detected in the presence of 4456dh (Fig. 5A). However, we observed a significant increase in trehalose content when *C. albicans* was exposed to high temperature stress (*P* < 0.0001; Fig. 5A). The addition of 0.5 mM 4456dh to *C. albicans* at 42 °C, resulted in an approximate two-fold decrease in trehalose content (Fig. 5A).

**Figure 5.**
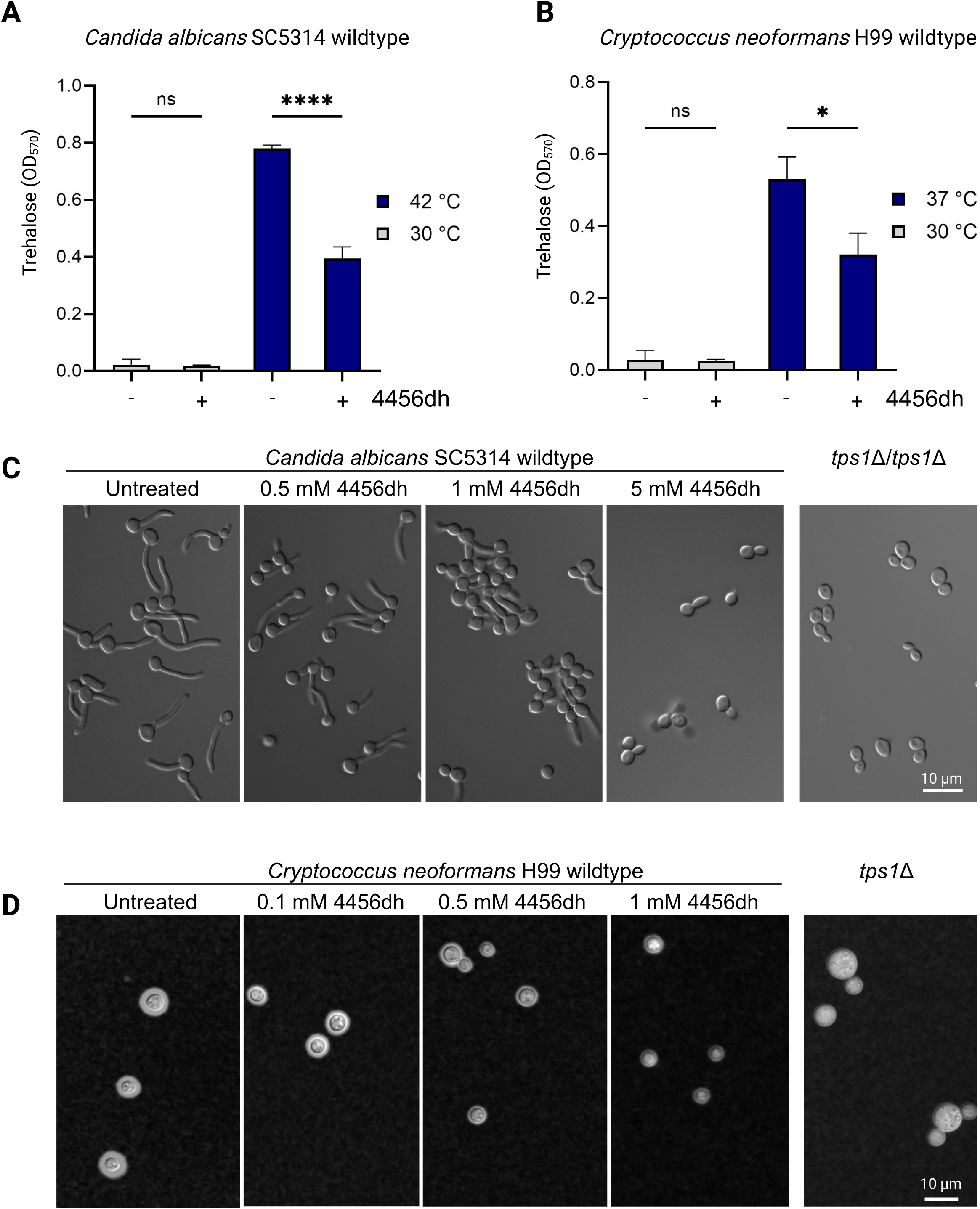
4456dh treatment results in reduced trehalose accumulation and a decrease in Tps1-dependent virulence factors. Trehalose accumulation measurements for *C. albicans* **(A)** and *C. neoformans* **(B)** cultured at indicated temperatures in the absence or presence of 0.5 mM 4456dh. Trehalose was measured using a colorimetric glucose assay after treatment of cultures with trehalase. Statistical comparisons were done with a one-way ANOVA analysis followed by a posthoc Tukey’s test. Error bars represent standard error. **C)** *C. albicans* wild-type strain SC5314 cells were treated with 0.5 mM, 1.0 mM and 5 mM 4456dh. Untreated *C. albicans* WT cells and the *C. albicans tps1*Δ/*tps1*Δ deletion strain were included in this experiment as positive and negative controls, respectively. Increasing concentrations of 4456dh negatively affect filamentation in *C. albicans* in YPD plus 10% fetal bovine serum incubated at 37 °C for 2 hours. Representative images from 2 independent experiments are shown. The scale bars represent 10 microns. **D)** Analysis of capsule formation in C. neoformans wild-type H99 using India Ink counter-staining in cells treated with 0.1, 0.5 mM and 1 mM 4456dh. Untreated *C. neoformans* WT cells and the *C. neoformans tps1*Δ deletion strain were included in this experiment as positive and negative controls, respectively. The cells were grown for 72 hours at 30 °C in capsule-inducing conditions. Cells were harvested, stained in India Ink and analyzed using differential interference contrast (DIC) imaging. Representative images from 3 independent experiments are shown. Scale bar represents 10 microns.

Similarly, we observed an increase in trehalose accumulation in *C. neoformans* cells grown at 37°C, compared to 30 °C (Fig. 5B). Treatment of *C. neoformans* with a sublethal dose of 4456dh at 37 °C resulted in a significant decrease in trehalose (*P* < 0.05; Fig. 5B). Furthermore, this 4456dh-mediated reduction in trehalose in *C. neoformans* occurred in a dose-dependent manner (Fig. S10).

Tps1 is required for serum-induced filamentation in *C. albicans* and production of polysaccharide capsule in *C. neoformans*, both of which are key virulence traits in these fungi (23, 28, 40). Therefore, we investigated whether 4456dh-mediated inhibition of Tps1 results in phenotypic recapitulation of a genetic deletion mutant lacking Tps1. Treatment of *C. albicans* cells with 4456dh during hyphal-inducing conditions resulted in defective filamentation in a dose-dependent manner with a complete inhibition of filamentation at sub-MIC concentration (5 µM) (Fig. 5C). Similarly, capsule production was affected in *C. neoformans* upon 4456dh treatment in a dose-dependent manner (Fig. 5D and S11). Combined, these assays directly show that 4456dh-mediated inhibition of Tps1 not only leads to less trehalose in fungal cells but also results in phenotypes that are hallmarks of lack of functional Tps1 in these fungi. These data are consistent with our hypothesis that 4456dh disrupts the substrate-binding capability of Tps1 and, therefore, significantly lowers trehalose biosynthesis (Fig. 6).

**Figure 6.**
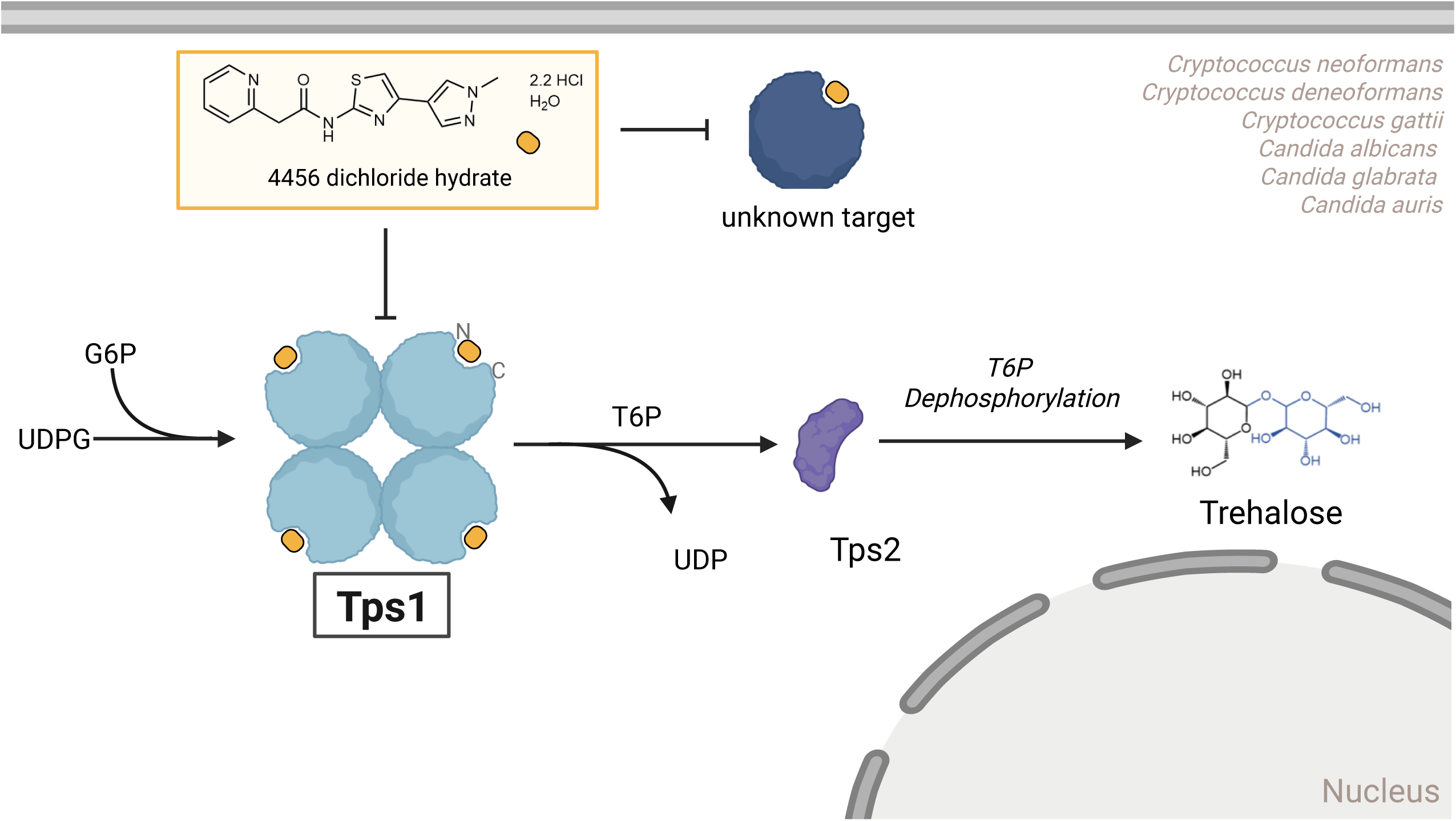
Model for 4456dh activity in fungal cells. Trehalose biosynthesis in *Cryptococcus* and *Candida* is initiated by Tps1, which converts G6P and UDPG into trehalose-6-phosphate (T6P). In the second step of the trehalose biosynthesis pathway, T6P is dephosphorylated by Tps2 and trehalose is generated. As shown, Tps1 from *C. neoformans* forms a homotetramer (30). 4456dh binds the exposed substrate-binding pockets of Tps1. Additionally, 4456dh may have an additional target that is likely another glycosyltransferase in some fungal organisms.

## Discussion

Fungal pathogens are devastating the ecosphere causing disease in agriculturally important crops, amphibians, bats and humans. These invasive fungal infections are prevalent worldwide and are responsible for millions of mortalities each year. To curb this significant rate of disease and death, the development of the antifungal arsenal needs to be continued and expanded to address concerns with antifungal drug resistance and the emergence of new fungal pathogens due to many factors, including a warmer climate.

Here we have focused on the development of a lead compound that targets Tps1 from *C. albicans* and *C. neoformans*, two predominant pathogenic fungal species in the clinics (1, 41). They have been highlighted as critical pathogens by the World Health Organization (3). Tps1 is a key enzyme in the critical first step of the trehalose biosynthesis pathway (15). Tps1 is a glycosyltransferase enzyme that uses substrates UDPG and G6P to generate the intermediate, trehalose-6-phosphate (T6P), which also acts as a signalling molecule. Tps1 is required for growth at 37 °C, capsule formation and growth on glucose in *C. neoformans* and *C. gattii* (24, 25, 40). *C. albicans* Tps1 also contributes to pathogenesis-related factors such as biofilm formation and hyphal transition from yeast (28). Perhaps most importantly, trehalose accumulates to high levels upon exposure of *Candida* and *Cryptococcus* to heat stress (24, 25). Tps1 is required for this response. Delineating this molecular mechanism and how to thwart it are critical to our complete understanding of how these fungal pathogens can survive at the elevated body temperature of the human host and cause disease.

4456dh is a soluble derivative of a lead compound identified in a screen to identify inhibitors of CaTps1 catalytic activity. We determined, using orthogonal methods, including microscale thermophoresis, thermal shift assays, a spectrophotometrically-coupled enzyme assay and x-ray crystallography, that 4456dh binds and competitively inhibits the activity of Tps1 and 4456dh has bioactivity against a broad range of *Candida* and *Cryptococcus* species (Fig. 6). Additionally, 4456dh also blocks the heat stress-induced hyperaccumulation of trehalose in both *C. albicans* and *C. neoformans* (Fig. 5).

We also discovered that 4456dh may have an additional target. Given that it is a small molecular weight compound, and GT-B glycosyltransferases have structurally conserved folds, it is not surprising. We hypothesize that 4456dh may target another glycosyltransferase and predict that it is one that does not involve a temperature-responsive phenotype because 4456dh retains partial activity at 30 °C in a *C. deneoformans tps1*Δ mutant. We identified multiple proteins in *C. albicans* and *C. neoformans* with structural similarity to Tps1 that may be affected by treatment with 4456dh (Fig. S12 and Tables S2 - 4). Future work will be needed to identify and characterize the additional target amongst these and other proteins.

There were great differences in the potency of 4456dh against fungal species. 4456dh minimal inhibitory concentration values were the lowest for *C. glabrata*, reaching 7.8 µM, whilst the other MICs were in the millimolar range for the fungi tested (Fig. 4). We were not able to find a clear structural explanation for the differences, given the significant conservation of the residues and structures of the substrate-binding pockets of Tps1 enzymes for these organisms (Fig. S5,6). Therefore, we hypothesize that the ability of 4456dh to enter the cell may underly the range of potency of 4456dh or the differences in the activity of 4456dh may also be due to the presence of a secondary protein target amongst the *Candida* and *Cryptococcus*. In the future, we aim to identify the additional target, or targets, of 4456dh using a directed, *in silico* approach to model 4456dh into glycosyltransferases in *Candida* and *Cryptococcus*. We shall also pursue a chemical genomics approach to determine targets of 4456dh.

Whilst previous studies found that *C. neoformans tps1*Δ cells were less susceptible to fluconazole at 30 °C, several studies have demonstrated that mutants in the trehalose biosynthesis pathway enhance susceptibility to antifungal drugs at elevated temperatures (25, 39, 42). Given that a proposed role of trehalose is to stabilize membranes during stress, such as elevated temperature, it is reasonable that loss of this protective sugar barrier would render the cell membrane more susceptible to attack by polyenes, azoles and echinocandins. Future studies will include investigations into the synergistic effects of 4456dh and the entire arsenal of the current antifungal therapeutics in several *Candida* and *Cryptococcus* species under stress.

Additional future structure-activity relationship studies using 4456dh will benefit from our initial structure of 4456dh bound to CaTps1. This scaffold, in addition to the previous structures of Tps1 will direct more readily the generation of new derivatives of 4456dh into a more effective Tps1-targeted drug. Our future studies will also include using medicinal chemistry to improve the selectivity and bioactivity of 4456dh. Although the MIC values of 4456dh are not like those of antifungals currently using in the clinic (Table S5), rational design of 4456dh analogues may increase the likelihood of using a 4456 derivative as an antifungal therapeutic in clinics. We predict that 4456dh analogues with increased specificity towards Tps1 will result in a compound with low toxicity because the trehalose biosynthesis pathway is not found in mammals.

In this study, we have provided proof of principle for successful pharmacological inhibition of Tps1, an enzyme in the trehalose biosynthesis pathway, which is central to fungal pathogen virulence and thermotolerance.

## Methods and Materials

### Strains

Fungal strains used in this study are listed in Supplemental Table 5. Strains were stored as glycerol stocks at −80 °C. Strains were grown at 30 °C on YPD (1% yeast extract, 2% Bacto Peptone, 2% dextrose) agar plates.

### Fluorescence polarization screen

The glycosyltransferase activity of 6xHis-CaTps1, which results in the release of UDP, was measured using the UDP Transcreener Fluorescence Polarization (FP) assay (BellBrook Labs #3018, Madison, WI). The St. Jude Children’s Hospital (Memphis, TN) 3-point pharmacophore (3PP) library was screened (36). Assays were performed at 37 °C for 60 minutes in a 384-well plate using 5 µM 6xHis-CaTps1 in assay buffer containing 50 mM Tris-HCl pH 7.5, 300 mM NaCl, 10 mM MgCl_2_ and 1 mM DTT. Fluorescence polarization was detected using a BMG Labtech plate reader. Each assay plate contained the following additional control wells: (i) no antibody control, (ii) DMSO vehicle control, (iii) novobiocin positive control and (iv) novobiocin dose-response. Screens were performed by addition of single point additions of test molecules to the above assay. The Z’ factor was calculated according to guidelines by Zhang *et*. *al*., with a Z-factor of 0.5 and 1.0 indicates an excellent assay (34).

### *De novo* co-crystallization and structure determination of *C. albicans* Tps1-4456 complex

*C. albicans* Tps1 (6xHis-CaTps1) in SEC buffer (20 mM Tris-HCl pH 8.0, 200 mM NaCl, 5% glycerol and 1 mM β-mercaptoethanol) was concentrated to 25 mg/mL using Amicon Ultra concentrators (30 MWCO, Millipore). 4456 which was added to the crystallization drop to a final concentration of 10 mM, from a 100 mM stock solution dissolved in 100% DMSO. Crystals were grown at 25 °C by hanging-drop vapor diffusion methods. Diffraction-quality crystals appeared after 3-4 days in a mother liquor solution containing 0.2 M magnesium sulfate, 0.1 M Tris-HCl pH 8.5 and 40% polyethylene glycol (PEG) 400. The crystals were cryo-preserved by looping and dipping a crystal for 1-2 seconds in a solution containing the crystallization reagent supplemented with 25% (v/v) glycerol. Data were collected at the Advanced Light Source (ALS) beamline 5.0.2 and processed with XDS (43). The structure of *C. albicans* Tps1-4456 was determined by molecular replacement using the *C. albicans* Tps1-UDP-G6P complex (PDB 5HUU, (28)) as the search model after removal of the UDP and G6P. Iterative cycles of model building were done in COOT (44) and refinement using Phenix refine. Selected data collection and refinement statistics are listed in Table 2.

Additional methodology for chemical synthesis, protein purification and several assays (binding, activity, MIC, fungicidal, trehalose, zone of inhibition, filamentation, E-tests and capsule) can be found in the Supplemental Material.

## Acknowledgements

E.J.W is supported by the Duke Science and Technology Initiative. This work was also funded by the grant NIH 1P01AI104533 to R.G.B., R.E.L., and J.R.P. J.R.P. is supported by grants AI73896 and A193257. V.Y. is supported by NIH/NIAID R01 awards AI039115-27 and AI172451-02 awarded to Joseph Heitman at Duke University Medical Center. C.B.N. is supported by grant funding from the US NIH R01 AI175711.

We thank all current and past Washington and Brennan laboratory members for helpful discussions. We are grateful to Dr. Joe Heitman for critical reading of the manuscript and Dr. Andrew Alspaugh for thoughtful discussions on this project. We thank Dr. Angela Rivera, Natalie Schulte and Dr. Peter Silinski for experimental support.

We thank the Advanced Light Source for access to 5.0.2. beamline. The Berkeley Center for Structural Biology is supported by the Howard Hughes Medical Institute, Participating Research Team members, and the National Institutes of Health, National Institute of General Medical Sciences, ALS-ENABLE grant P30 GM124169. The Advanced Light Source is a Department of Energy Office of Science User Facility under Contract No. DE-AC02-05CH11231.

Y.M., V.Y., W.S. and E.J.W. conducted most experiments, and interpreted data and results. J.L., A.J., C.B.N., C.G., J.L.T. and M.S. conducted some experiments and interpreted data and results. E.J.W. wrote the original draft of the manuscript. E.J.W., R.E.L., R.G.B and J.R.P. conceived the study and interpreted data and results. E.J.W., V.Y., R.E.L., R.G.B and J.R.P. assisted with editing the manuscript.

## Data Availability

Atomic models of CaTps1-SJ6675 and CaTps1-4456 have been deposited in the RCSB Protein Data Bank (PDB) (https://rcsb.org) with accession codes PDB ID 9NKB and PDB ID 9NIQ, respectively. All additional data, including raw data and images associated with all figures, is available upon reasonable request to R.G.B. and E.J.W., the corresponding authors.

