## Supplemental Material for "Inhibitors of trehalose-6-phosphate synthase activity in fungal pathogens compromise thermal tolerance pathways"

[1] co-first authors

### Supplemental Methods and Materials

**Synthesis of N-(4-(1-methyl-1H-pyrazol-4-yl)thiazol-2-yl)-2-(pyridin-2-yl)acetamide Dichloride Hydrate(4456).**

To a stirred solution of 4-(1-methyl-1H-pyrazol-4-yl)thiazol-2-amine (**1,** 1.75 g, 9.71 mmol) in dry DMF (20mL) at room temperature, a single portion of 2-(pyridin-2-yl)acetic acid hydrochloride (**2**, 3.71 g, 21.36 mmol) and HBTU (8.10 g, 21.36 mmol) was added. The resulting mixture was stirred at room temperature for 5 minutes, followed by the addition of DIPEA (5.59 g, 48.50 mmol). The reaction was stirred for 1 hour before being partitioned between dichloromethane and water. The organic layer was washed with brine, dried over $Na_2SO_4$ and the solvent was removed under reduced pressure. The residue was redissolved in ethyl acetate (EA), and a pale solid was obtained by filtration (1.25g, 43.0%). To convert the product to its dichloride form, 20 mL of 1M HCl in MeOH was added to the suspended solution of **4456** (1.25g) in MeOH. Once the suspending solution became clear, the solvent was evaporated, and the residue was treated with EA. The solid was collected by filtration as **4456** dichloride hydrate (1.66g, 100%). $^1$H NMR (400 MHz, $D_2O$) δ 8.81 (dd, J = 6.3, 1.7 Hz, 1H), 8.61 (td, J = 8.0, 1.6 Hz, 1H), 8.12 – 7.87 (m, 4H), 7.18 (s, 1H), 3.92 (s, 3H), 3.35 (s, 2H). Anal. Calcd for $C_{14}H_{13}N_5OS$ $H_2O$ 2.2HCl: C 42.29, H 4.36, N 17.62, Cl 19.62; Found C 42.52, H 4.53, N 17.33, Cl 19.46.

**Chemistry**

4-(1-methyl-1H-pyrazol-4-yl)thiazol-2-amine (**1**) was purchased from Enamine. $^1$H NMR spectra were recorded on a Bruker 400 MHz NMR spectrometer. Chemical shifts (δ) are reported in parts per million relative to the residual solvent peak or internal standard (tetramethylsilane) and coupling constants (J) are reported in hertz (Hz). The purity of the products was confirmed by UPLC/MS (the Waters Acquity). Elemental analysis was tested by Atlantic Microlab Inc.

**Expression and Purification of *C. albicans* Tps1**

*C. albicans* Tps1 (CaTps1) was purified as previously described (1). Briefly, the full-length *TPS1* gene from *C. albicans* strain SC5314 was codon-optimized for expression in *E. coli* (Genscript) and cloned into the pET-28a vector, which contains an N-terminal 6xHis affinity tag followed by a thrombin cleavage site. The construct was transformed into BL21(DE3)pLysS cells (Life Technologies, Inc.) and induced with 0.5 mM isopropyl β-D-1-thiogalactopyranoside (IPTG) at 15 °C overnight. Supernatants from lysed cultures were loaded onto a nickel column (Ni-NTA, Qiagen) and washed in a buffer containing 50 mM Tris-HCl pH 8.0, 300 mM NaCl, 5 mM $MgCl_2$, 5% glycerol and 5 mM imidazole. The protein was eluted with increasing amounts of imidazole in the wash buffer. The fractions containing 6xHis-CaTps1 were pooled, reduced to 5 mL and purified further using S200 size exclusion column chromatography (HiLoad 26/600 Superdex 200pg, Cytiva) in a precooled buffer containing 20 mM Tris-HCl pH 8.0, 200 mM NaCl, 5% glycerol and 1 mM β-mercaptoethanol. 5 mL fractions from the size exclusion column containing 6xHis-CaTps1, as determined by SDS-PAGE analysis, were pooled and concentrated using 30K MWCO Amicon Ultra concentrators (Millipore) to 1 mg/mL for downstream applications.

**Expression and Purification of *C. neoformans* Tps1**

*C. neoformans* Tps1 (CnTps1) was purified as previously described (2). Briefly, the full-length *TPS1* gene from *C. neoformans* strain H99 was codon-optimized for expression in *E. coli* (Genscript) and subcloned using ligation-independent cloning into pMCSG7 (3). The construct was transformed into *E. coli* OverExpress C41(DE3) chemically competent cells engineered for high protein expression (Sigma). Supernatants from lysed cultures, induced with 0.5 mM isopropyl β-D-1-thiogalactopyranoside (IPTG), were loaded onto a nickel column (Ni-NTA, Qiagen) and washed in a buffer containing 50 mM Tris-HCl pH 8.0, 300 mM NaCl, 5 mM $MgCl_2$, 5% glycerol and 5 mM imidazole. The protein was eluted with increasing amounts of imidazole in the wash buffer. The fractions containing 6xHis-CnTps1 were pooled, reduced to 5 mL and purified further using S200 size exclusion column chromatography (HiLoad 26/600 Superdex 200pg, Cytiva) using a precooled buffer containing 20 mM Tris-HCl pH 8.0, 300 mM NaCl, 5% glycerol and 2 mM β-mercaptoethanol. 5 mL fractions from the size exclusion column containing 6xHis-CnTps1, as determined by SDS-PAGE analysis, were pooled and concentrated to 1 mg/mL for downstream applications.

**Microscale Thermophoresis Assays**

Recombinant and 6xHis-tagged CaTps1 and CnTps1 were purified according to the procedures described above. CaTps1 and CnTps1 were diluted to 5 µM in buffer containing (SEC buffer with 0.05% Tween20). For the dilution series, a 100 µM solution of 4456dh was prepared using the same buffer. The stock solution was used for the 16-step serial dilution in the buffer, with a final volume of 10 µL of 4456dh in each reaction mixture of the dilution series. 10 µL of protein was added to the 16 vials and samples were mixed by pipetting up and down. Reactions were incubated overnight at room temperature. Samples were loaded into Monolith NT LabelFree Premium capillaries. MST was performed at Medium MST power for both CaTps1 and CnTps1. The data were acquired with MO.Control 1.5.3 (NanoTemper Technologies GmbH). Recorded data were analyzed with MO.Affinity Analysis 2.2.7 (NanoTemper Technologies GmbH). The MST on-time that yielded that highest signal-to-noise ratio was used for the $K_d$ determination, where $K_d$ is the equilibrium dissociation constant.

**Thermal Shift Assays**

6xHis-CaTps1 and 6xHis-CnTps1 (2 µM) were individually incubated with 4456dh (100 µM) in 20 mM Tris-HCl pH 8.0, 300 mM NaCl, 5% glycerol and 2 mM β-mercaptoethanol buffer containing 2X Glo-melt dye. Thermal denaturation was measured using an RT-PCR thermocycler (BioRad) between 25 °C and 95 °C at 0.5 °C increments. The melting temperatures of each sample were determined by identifying the inflection point of the derivative data.

**Tps1 Enzyme Activity Assay**

The catalytic activity of *C. albicans* Tps1 and *C. neoformans* Tps1 was measured via a continuous enzyme-coupled assay as previously reported (4). Briefly, either 6xHis-CaTps1 or 6xHis-CnTps1 were concentrated in a buffer containing 20 mM Tris-HCl pH 8.0, 300 mM NaCl, 5% glycerol and 2 mM β-mercaptoethanol. The assay was performed in an assay buffer containing 50 mM HEPES pH 7.8, 100 mM KCl, 5 mM $MgCl_2$ and 2 mM DTT. Final concentrations of 3 µM protein were combined with 1 mM UDPG and 1 mM G6P. Activity assays were performed in clear, flat-bottomed 96-well microtiter plates and the decrease in absorbance at 340 nm during the initial reaction rate was recorded using a plate reader (Tecan). The decrease in absorbance was analyzed for the first 200s of the kinetic reaction.

**Minimum Inhibitory Concentration Assay**

Wild-type *C. albicans* strains SC5314, wild-type *C. glabrata* CBS138 and wild-type *C. auris* B11220 were cultured overnight in YPD (yeast-peptone-dextrose) medium at 30 °C. Growth of the overnight cultures was quantified by measuring the $OD_{600}$ and corrected for the background absorbance of the medium. Antifungal potency for 4456dh was assessed by minimum inhibitory concentration (MIC) dose-response assays performed under standard Clinical and Lab Standards Institute (CLSI) conditions. A volume of 125 µL from each diluted cell suspension was dispensed into the wells of 96-well flat-bottom microtiter plates (Corning). An additional 25 µL of 4456dh from a series of dilutions from the 157 mM stock solution prepared in water was added to each well. Following preparation, the plates were incubated at the either 30 °C or 42 °C, as indicated, for 48 hours. All dose-response assays were performed in biological triplicate with a minimum of three technical replicates. Growth, as determined by a reading of absorbance at 600 nm, was normalized to untreated controls and was corrected for the absorbance of 4456dh in the YPD medium. Growth was plotted as a heat map using Excel.

Wild-type *C. neoformans* strains H99, wild-type *C. deneoformans* JEC21 and wild-type *C. gattii* R265 were cultured overnight in YPD (yeast-peptone-dextrose) medium at 30 °C. Growth of the overnight cultures was quantified by measuring the $OD_{600}$ and corrected for the background absorbance of the medium. Antifungal potency for 4456dh was assessed by minimum inhibitory concentration (MIC) dose-response assays performed under standard Clinical and Lab Standards Institute (CLSI) conditions. A volume of 125 µL from each cell suspension was dispensed into the wells of 96-well flat-bottom microtiter plates (Corning). An additional 25 µL of 4456dh from a series of dilutions from the 157 mM stock solution prepared in water was added to each well. Following preparation, the plates were incubated at the either 30 °C or 37 °C, as indicated, for 72 hours.

All dose-response assays were performed in biological triplicate with a minimum of three technical replicates. Growth, as determined by a reading of absorbance at 600 nm, was normalized to untreated controls and was corrected for the absorbance of 4456dh in the YPD medium. The minimum inhibitory concentration causing 80% growth reduction, or $MIC_{80}$, was determined based on the $OD_{600}$ values (5, 6). Growth was plotted as a heat map using Excel.

**Fungicidal Assays**

Cells previously exposed to compounds for 48 or 72 hours in dose-response matrices were spotted to test the viability of treated cells. 3 µL was extracted from each well with a multi-channel pipette and dispensed onto drug-free YPD agar plates. Plates were incubated at 30 °C for 48 hours and photographed. All experiments were performed in biological and technical duplicate.

**Trehalose Measurement**

*Candida* and *Cryptococcus* strains were grown in liquid cultures at 30 °C and then 42 °C or 37 °C, respectively. Cells at each temperature were exposed to either water (vehicle control) or 0.5 mM 4456dh. The $OD_{600}$ was determined followed by dilution of overnight cultures to an equivalent of 0.1 $OD_{600}$ of cells. Cells were pelleted, frozen and lyophilized and stored in -80 °C. Pellets were lysed by vortexing with sterile glass beads. The cell-free extract was generated by adding 1X PBS and centrifuging the tubes to pellet the glass beads. The supernatant containing cell-free extracts was exposed to trehalase (Sigma) overnight at 37 °C and then tested for trehalose levels according to the Glucose Assay Kit (MAK476, Sigma) protocol.

**Zone of Inhibition Assays**

Strains were grown in overnight cultures in YPD (yeast-peptone-dextrose) medium at 30 °C. Optical density ($OD_{600}$) was determined with an Infinite PRO plate reader (Tecan). $OD_{600}$ was adjusted to 0.01 ($10^6$ cells/mL) through dilution with YPD, and 200 µL diluted culture from each strain was plated onto YPD plates and spread using sterile beads (Sigma). After the plates had dried, a single disk soaked with either 15 µL of water, 4456dh or 1 µg/mL FK506 disk (6 mm diameter, Becton, Dickinson and Company) was placed in a quadrant of each plate. FK506 was used as the positive control (7). Plates were incubated at either 30 °C or 37 °C for 48 h and then imaged.

***C. albicans* Filamentation Assays**

*C. albicans* wild-type strain SC5314 and respective *tps1*/*tps1* deletion strains were grown overnight at 30°C in 5 ml liquid YPD. 1 OD equivalent of the overnight grown cells were harvested, resuspended in 1 ml of YPD+10% fetal bovine serum and incubated at 37°C for 2 hours. The wild-type culture was grown in four replicates and three of them were treated with Lee4456dh at different concentrations (0.5mM, 1mM, and 5mM) and one was used as untreated control. *tps1*/*tps1* deletion mutant was used as a negative control for filamentation in these assays. 10 µl of cell suspension was directly used for imaging the filamentation status of the cells using the Zeiss Axio Scope microscope attached to the Axiocam at 40X magnification. The images were processed using Fiji (8) for the final presentation.

**Epsilometer Test (E-test)**

*C. neoformans* wild type (H99) was incubated in a shaking culture of YPD medium for 18 h at 30ºC. The culture was diluted to an $OD_{600}$ 0.6, diluted 10-fold in PBS, and 100 µL was spread onto 60 X 15 mm YPD agar plates prepared with different concentrations of 4456dh. After the surface was completely dry, an E-test strip containing a gradient of fluconazole (Biomérieux, Marcy-l'Étoile, France) was applied to each plate. The plates were incubated at 37 °C and imaged after 72 h.

***C. neoformans* Capsule Assays**

Capsule was induced by inoculating fresh *C. neoformans* wild-type strain H99 cells in $CO_2$-independent liquid media (Gibco) for 72 hours at 30 °C. The wild-type culture was grown in four replicates and three of them were treated with Lee4456dh at different concentrations (0.1 mM, 0.5 mM and 1 mM) and one was used as untreated control. The *C. neoformans tps1*Δ mutant was used as a negative control for capsule formation. For visualization, BactiDrop India Ink (Remel) was added to the cell suspension and 5 µL of this mixture was then spotted onto a glass slide. Capsule was observed with a Zeiss Axio Imager A1 fluorescence microscope with a 100X objective. Images were taken with an AxioCam MRm digital camera with ZEN Pro software (Zeiss). The size of the capsule of > 100 cells per 4456dh treatment was quantified using Image J software (https://imagej.net/ij/index.html) by measuring the width of the halo created by capsule and made visible with the negative India Ink staining.

### Supplemental Figure Legends

**Supplemental Figure 1. Fluorescence polarization dose-response curves.** Fluorescence polarization binding curves of hits from the 3-point pharmacophore (3PP) library. Two wells per data point. Error bars represent standard error.

**Supplemental Figure 2. A structure-guided approach led to the derivatization of SJ6675**. **A)** CaTps1-SJ6675 crystal structure at resolution of 3.5 Å. The zoomed-in view shows SJ6675 in the proximity of CaTps1 substrate-binding residues R280, K285, N382 and L383. **B)** The chemical structures and identities of the SJ6675 derivative library. 4456, the focus of additional studies presented here, is highlighted. **C)** Effect of derivatives on the activity of 6xHis-CaTps1. CaTps1 enzymatic activity was tested using a coupled activity assay and normalized to the DMSO control (data represent the mean ± SEM, n = 3). Compounds that precipitated are represented with the grey bars. The compounds that did not precipitate and the DMSO vehicle control are represented as purple and blue, respectively.

**Supplemental Figure 3. 4456dh NMR validation and elemental analysis. A)** $^{1}$H NMR spectra were recorded on a Bruker 400 MHz spectrometer. Chemical shifts (δ) are reported in parts per million relative to the residual solvent peak or internal standard. Coupling constants (J) are reported in hertz (Hz). **B)** Elemental analysis confirmed the chemical composition of C, H, N and Cl elements in 4456dh.

**Supplemental Figure 4. Polder omit map of 4456 in the CaTps1-4456 structure**. A polder omit map**,** contoured at 3σ (purple mesh), shows 4456 electron density bound in the CaTps1 substrate-binding site. CaTps1 residues and 4456 are shown as atom-colored stick and ball representations.

**Supplemental Figure 5. Residues within 4 Å of 4456 in the *C. albicans* Tps1-4456 complex are conserved.** Tps1 sequences of *C. gattii*, *C. deneoformans*, *C. neoformans*, *A. fumigatus*, *C. glabrata*, *C. auris* and *C. albicans* are aligned. The sequences are taken from the NCBI database. Key residues within 4 Å of 4456 are highlighted by red boxes. The numbering in the red text is based on the *C. albicans* Tps1 protein sequence.

**Supplemental Figure 6. 4456 binds in the structurally conserved pocket of Tps1 enzymes. A)** Structural alignments of one subunit of the Tps1 enzymes from *C. albicans* (pink, PDB: 5HUU), *E. coli* (purple, 1UQU), *C. neoformans* (green, 8FHW) and *A. fumigatus* (blue, 5HVM) with 4456 (sand) from the *C. albicans* Tps1-4456 complex modelled into the conserved catalytic pocket of Tps1 (1, 2, 9). **B)** A ConSurf analysis for CaTps1 and 4456 (as a grey, surface-representation). The amino acids are colored by their levels of conservation as shown with the color-coding bar. Teal represents amino acids that are variable, whereas dark magenta amino acids are highly conserved. Regions for which there are not enough data to form a conclusion are shown in yellow.

**Supplemental Figure 7. 4456dh bioactivity at permissive growth temperatures (30 °C) for *Candida* and *Cryptococcus*. A)** Table showing minimum inhibitory concentration values for 4456dh in *Candida* and *Cryptococcus* species grown at 30 °C. **B)** *Candida* spot assays show ability of cells to grow after exposure to 4456 and indicate fungicidal activity of 4456dh at 30 °C. **C)** *Cryptococcus* spot assays show ability of cells to grow after exposure to 4456 and indicate fungicidal activity of 4456dh at 30 °C.

**Supplemental Figure 8. 4456dh retains activity in a *C. deneoformans tps1*Δ. A)** Zone of inhibition assays with *C. deneoformans* strains JEC21 and JEC20 grown at 37 °C. **B)** Zone of inhibition assays with *C. deneoformans* strains JEC21 *tps1*Δ and JEC20 *tps1*Δ grown at 30 °C.

The circular graphic illustrates the compounds added to the sterile disc. FK506 (1 μg/mL), a calcineurin inhibitor, is the positive control and water is the vehicle control.

**Supplemental Figure 9. Synergy between 4456dh and fluconazole in *C. neoformans*.** Images of YPD plates containing serially-diluted concentrations of 4456dh with fluconazole E-test strips. The plate lacking 4456dh is the negative control. Zones of *C. neoformans* growth inhibition indicate the relative susceptibility of the fungi to the combination of drugs.

**Supplemental Figure 10. 4456dh dose-dependent decrease in trehalose in *C. neoformans*.** Trehalose accumulation measurements for *C. neoformans* cultured at 37 °C in the presence of 0.1, 0.5, 1.0 and 2.0 mM 4456dh. The untreated sample served as the negative control. Trehalose was measured using a colorimetric glucose assay after treatment of cultures with trehalase. Error bars represent standard error.

**Supplemental Figure 11. Quantification of capsule formation in *C. neoformans* treated with 4456dh.** Images taken were analyzed with ImageJ. A minimum of 100 cells per treatment or strain were analyzed. The median for each sample is indicated with the black line. Statistical analysis was performed in GraphPad Prism using the one-way ANOVA with Bartlett's posthoc test to compare the untreated sample to the individual treated samples. P values $< 0.05$ are indicated as significant on the graph.

**Supplemental Figure 12. Selected predicted structures with structural homology to *C. albicans* Tps1.** The query *C. albicans* Tps1 (PDB: 5HUU, Chain A) is shown in grey (1). AlphaFold predicted structures, identified by a hierarchical search of the DALI protein structure comparison server, are shown in ribbon representations and colored according to the pLDDT (predicted local distance difference test) (10).

### Supplemental Tables

**Supplemental Table 1. Data collection and refinement statistics of the structure of 6xHis-CaTps1 in complex with compound SJ6675.**

| **6xHis-CaTps1 - SJ6675 complex** | |
|---|---|
| Resolution range | 47.05 - 3.501 (3.63 - 3.5) |
| Space group | P $6_5$ 2 2 |
| Unit cell | 115.1 115.1 266.8 90 90 120 |
| Total reflections | 157,931 (14,753) |
| Unique reflections | 13,858 (1,326) |
| Multiplicity | 11.4 (11.1) |
| Completeness (%) | 99.60 (99.10) |
| Mean I/sigma(I) | 17.15 (6.63) |
| Wilson B-factor | 94.6 |
| R-merge | 0.0969 (0.2914) |
| R-meas | 0.1017 (0.306) |
| R-pim | 0.03055 (0.09207) |
| $CC_{1/2}$ | 0.999 (0.971) |
| CC* | 1 (0.993) |
| Reflections used in refinement | 13,814 (1324) |
| Reflections used for R-free | 1,366 (133) |
| R-work | 0.2300 (0.3096) |
| R-free | 0.2748 (0.3868) |
| Number of non-hydrogen atoms | 7,474 |
| macromolecules | 7,474 |
| ligands | 0 |
| solvent | 0 |
| Protein residues | 934 |
| RMS(bonds) | 0.003 |
| RMS(angles) | 0.64 |
| Ramachandran favored (%) | 96.34 |
| Ramachandran allowed (%) | 3.23 |
| Ramachandran outliers (%) | 0.43 |
| Rotamer outliers (%) | 0.00 |
| Clashscore | 6.65 |
| Average B-factor | 92.46 |
| macromolecules | 92.46 |

Statistics for the highest-resolution shell are shown in parentheses.

**Supplemental Table 2. Similarity between *C. albicans* Tps1 and Alphafold predicted glycosyltransferase structures in *C. albicans*.**

| AF# | Z-score | RMSD | %ID | Protein |
|---|---|---|---|---|
| Q92410 | 47.8 | 1.1 | 64 | Alpha, Alpha-trehalose phosphate synthase |
| A0A1D8PIS4 | 38.5 | 1.4 | 32 | Trehalose 6-phosphate synthase/phosphatase |
| Q5AI14 | 37.0 | 1.7 | 35 | Trehalose-phosphatase |
| Q59LF2 | 25.0 | 2.9 | 13 | Alpha-1,3/1,6-mannosyltransferase Alg2 |
| Q59Q79 | 21.8 | 3.6 | 11 | Chitobiosyldiphosphodolichol beta mannosyltransferase |
| Q5A6R7 | 20.6 | 3.2 | 13 | Glcnac-Pi Synthesis protein |
| Q59S72 | 16.3 | 3.2 | 9 | GDP-Man:Man(3)GlcNAc(2)-PP-Dol alpha-1,2-mannosyltransferase Alg11 |
| Q5A850 | 16.0 | 4.0 | 11 | Glycogen synthase |
| A0A1D8PQQ3 | 13.8 | 3.7 | 9 | Alpha-1,4 glucan phosphorylase |
| Q5A950 | 9.6 | 4.4 | 7 | Sterol 3-beta-glucosyltransferase Atg26 |
| Q5A868 | 8.5 | 4.6 | 9 | Phosphopantothenoylcysteine decarboxylase complex subunit |
| Q5ABE5 | 8.3 | 4.6 | 10 | UDP-N-acetylglucosamine transferase subunit ALG13 |
| A0A1D8PQJ9 | 7.8 | 3.1 | 10 | Ubiquinone biosynthesis protein |
| Q5A1C0 | 7.7 | 3.1 | 9 | Uncharacterized protein |
| Q5AK46 | 7.5 | 4.2 | 6 | Multifunctional fusion protein |
| Q5AML3 | 7.5 | 3.2 | 10 | Oxidoreductase |
| A0A1D8PE77 | 7.4 | 4.8 | 7 | Ubiquitin-activating enzyme E1-like |
| A0A1D8PKD2 | 7.3 | 5.2 | 7 | Phosphopantothenoylcysteine decarboxylase complex subunit |
| A0A1D8PKJ3 | 7.3 | 10.3 | 3 | E1 ubiquitin-activating protein UBA1 |
| Q59XU5 | 7.2 | 3.4 | 6 | Ras-like protein 1 |
| A0A1D8PCN0 | 7.0 | 3.4 | 8 | Bifunctional UDP-glucose 4-epimerase/aldose 1-epimerase |
| Q5AP65 | 7.0 | 3.4 | 11 | Protein FMP52, mitochondrial |
| Q5A5N6 | 7.0 | 10.0 | 12 | UDP-N-acetylglucosamine transferase subunit ALG14 |
| A0A1D8PKJ4 | 6.9 | 3.1 | 13 | Saccharopine dehydrogenase (NADP+, L-glutamate-forming) |
| A0A1D8PRL3 | 6.9 | 3.5 | 10 | NAD(P)-bd domain-containing protein |

A hierarchical search for query structure against the AlphaFold database using the DALI protein structure comparison server. The query *C. albicans* Tps1 (PDB:5HUU, Chain A) was used to search for similar predicted structures generated by AlphaFold in *C. albicans* (1).

**Supplemental Table 3. Structures in the PDB that have similarity to *C. neoformans* Tps1 (PDB: 8FO1).**

| PDB | RMSD (Å) | %ID | Aligned Residues | Protein | Organism |
|---|---|---|---|---|---|
| 6PTA | 2.43 | 5 | 94 | ARF family small GTPase ARF1 in complex with GDP | *C. albicans* |
| 7RLL | 2.46 | 10 | 93 | ARF3 in complex with guanosine-3'-monophosphate-5'-diphosphate | *C. albicans* |
| 8PHX | 2.67 | 9 | 79 | Receiver Domain of the Hybrid Histidine Kinase Sln1 | *C. albicans* |
| 5UF8 | 2.71 | 7 | 95 | ARF family small GTPase ARF2 in complex with GDP | *C. albicans* |
| 6EOA | 3.23 | 10 | 125 | Crystal Structure of HAL3 | *C. neoformans* |
| 6EFW | 3.28 | 6 | 119 | YjeF family protein | *C. neoformans* |
| 6EFX | 3.58 | 10 | 130 | YjeF family protein in complex with AMPPNP | *C. neoformans* |
| 8JAT | 2.52 | 11 | 95 | 3-ketodihydrosphingosine reductase TSC10 | *C. neoformans* |

A search for query structure against the NCBI VAST+ Similar Structures search algorithm. The query *C. neoformans* Tps1 (PDB: 8FO1) was used to search for similar predicted structures found in the PDB (2).

**Supplemental Table 4. Structures in the PDB that have similarity to *C. albicans* Tps1 (PDB: 5HUU).**

| PDB | RMSD (Å) | %ID | Aligned Residues | Protein | Organism |
|---|---|---|---|---|---|
| 7RLL | 2.52 | 7 | 97 | ARF3 in complex with guanosine-3'-monophosphate-5'-diphosphate | *C. albicans* |
| 7DLL | 2.41 | 7 | 87 | Short chain dehydrogenase 2 (SCR2) with NADPH | *C. parapsilosis* |
| 7DMG | 3.00 | 6 | 97 | Short chain dehydrogenase 2 (SCR2) with NADP | *C. parapsilosis* |
| 7VYQ | 3.11 | 5 | 104 | Short chain dehydrogenase (SCR) with NADP and ethyl 4-chloroacetoacetate | *C. parapsilosis* |
| 7DLD | 3.13 | 12 | 109 | (S)-carbonyl reductases in different oligomerization states | *C. parapsilosis* |
| 7DLM | 3.20 | 7 | 104 | Short chain dehydrogenase (SCR) with NADPH | *C. parapsilosis* |
| 8JAT | 2.65 | 12 | 98 | 3-ketodihydrosphingosine reductase TSC10 | *C. deneoformans* |

A search for query structure against the NCBI VAST+ Similar Structures search algorithm. The query *C. albicans* Tps1 (PDB: 5HUU) was used to search for similar predicted structures found in the PDB (1).

**Supplemental Table 5. Comparison of Minimum Inhibitory Concentrations of 4456dh and clinically relevant antifungal therapeutics.**

| Drug | Pathogen | $MIC_{80}$ (µg/mL) | Reference |
|---|---|---|---|
| 4456dh | *C. albicans* | 5160 | This publication |
| 4456dh | *C. auris* | 2780 | This publication |
| 4456dh | *C. glabrata* | 3.1 | This publication |
| 4456dh | *C. neoformans* | 2780 | This publication |
| 4456dh | *C. deneoformans* | 1190 | This publication |
| 4456dh | *C. gattii* | 2780 | This publication |
| Fluconazole | C. albicans | 1.84 | (11) |
| Voriconazole | C. albicans | 0.06 | (11) |
| Caspofungin | C. albicans | 1 | (12) |
| Amphotericin B | C. neoformans | 2 | (13) |
| Flucytosine | C. neoformans | 8 | (13) |

**Supplemental Table 6. Fungal strains used in this study.**

| Strain Name | Genotype | Source |
|---|---|---|
| *C. albicans* SC5314 | Wild-type | (14) |
| *C. auris* B11220 | AR0381 | (15) |
| *C. glabrata* CBS138 | Wild-type | (16) |
| *C. neoformans* H99α | Wild-type *MAT*α | (17) |
| *C. deneoformans* JEC20 | Wild-type *MATa* | (18) |
| *C. deneoformans* JEC21 | Wild-type *MAT*α | (18) |
| *C. gattii* R265 | *C. gattii* wild-type VGII mating type α | (19) |
| *C. deneoformans* JEC20 *tps1*Δ | MATa*tps1*::*NAT* | (20) |
| *C. deneoformans* JEC21 *tps1*Δ | MATα *tps1*::*NAT* | (20) |

A

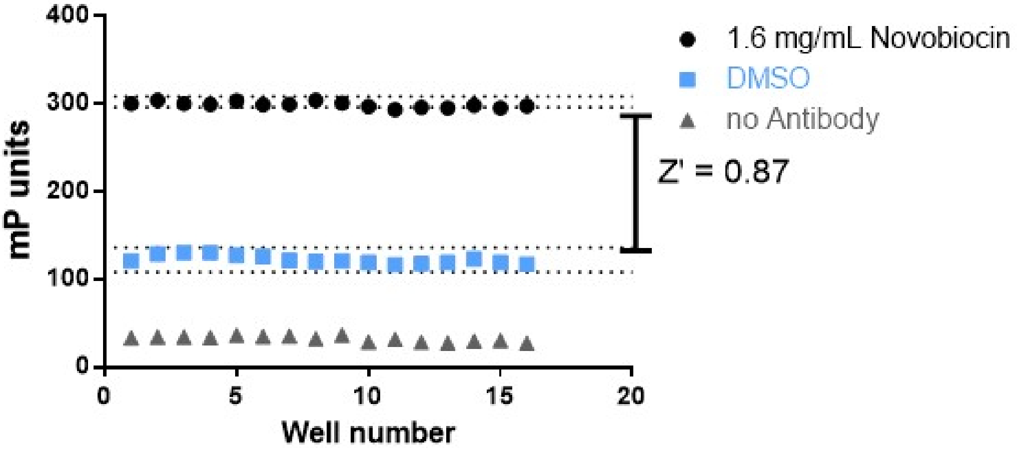
400
300
200
100
0
mP units
0
5
10
15
20
Well number
1.6 mg/mL Novobiocin
DMSO
no Antibody
Z' = 0.87

B

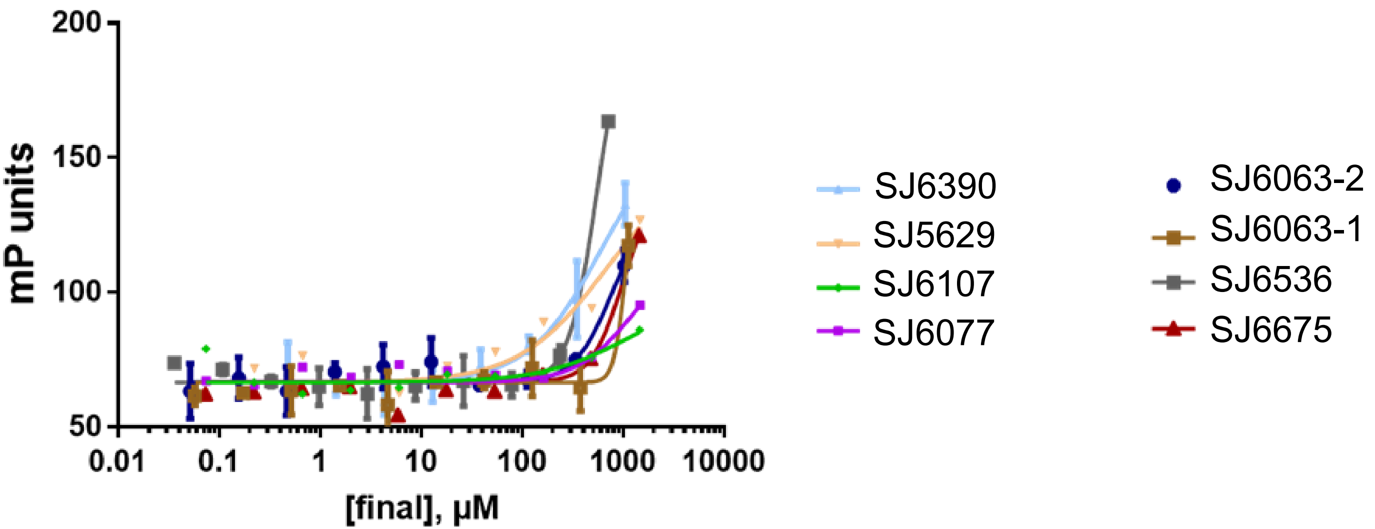
200
150
100
50
mP units
0.01
0.1
1
10
100
1000
10000
[final], µM
SJ6390
SJ5629
SJ6107
SJ6077
SJ6063-2
SJ6063-1
SJ6536
SJ6675

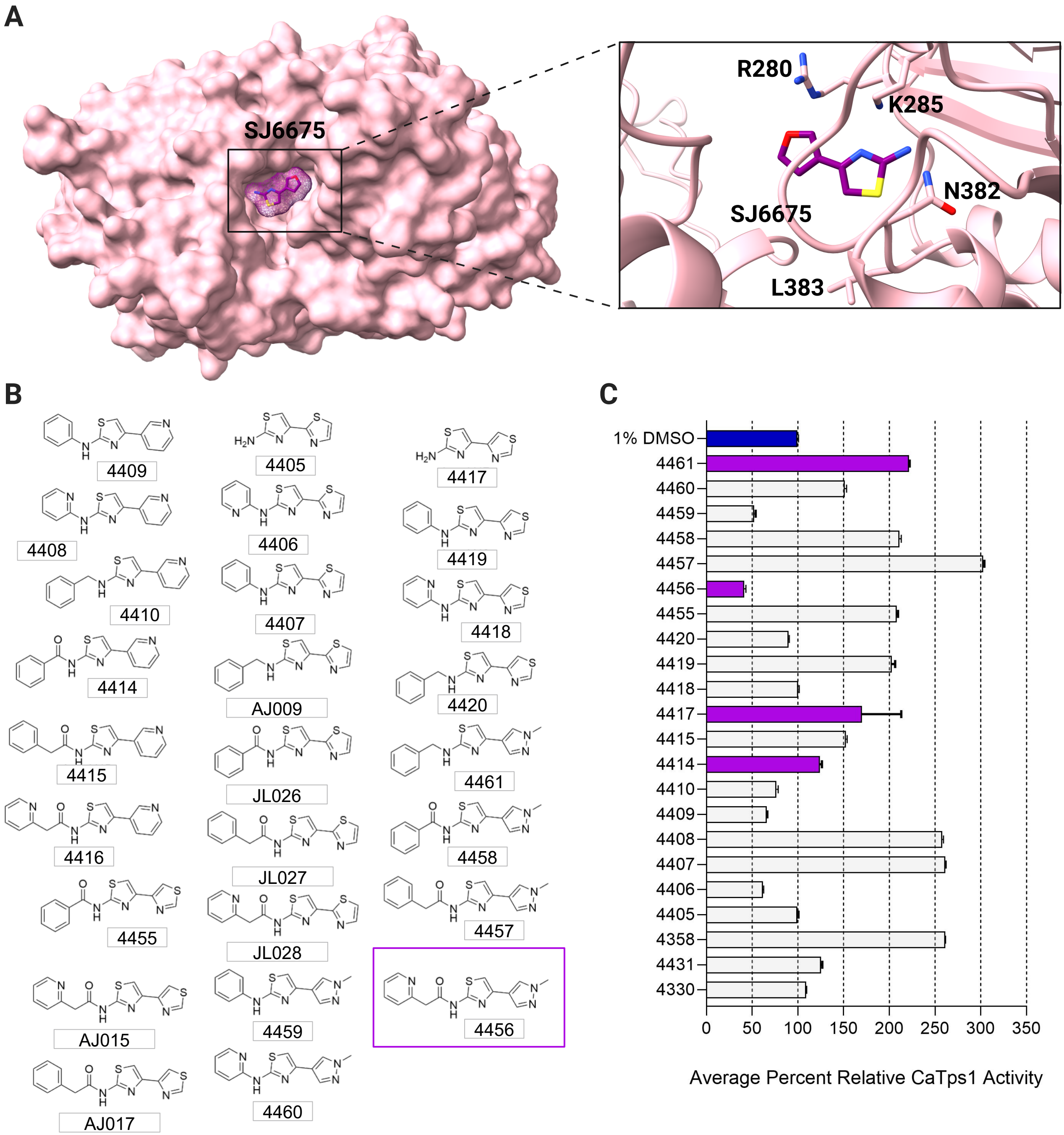
A
SJ6675
R280
K285
N382
SJ6675
L383
B
4409
4408
4410
4414
4415
4416
4455
AJ015
AJ017
4405
4406
4407
AJ009
JL026
JL027
JL028
4459
4460
4417
4419
4418
4420
4461
4458
4457
4456
C
1% DMSO
4461
4460
4459
4458
4457
4456
4455
4420
4419
4418
4417
4415
4414
4410
4409
4408
4407
4406
4405
4358
4431
4330
0
50
100
150
200
250
300
350
Average Percent Relative CaTps1 Activity

**A**

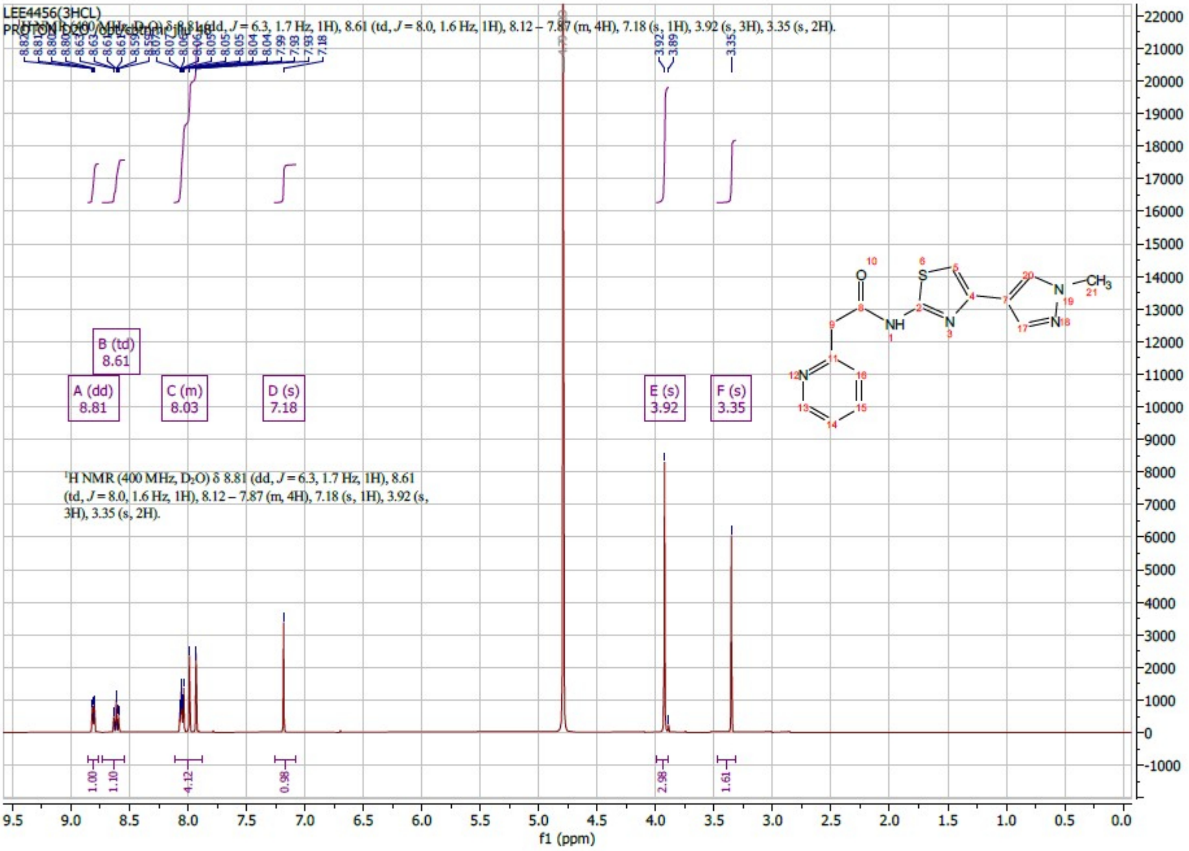


**B**

| Element | Theory | Found | |
|---|---|---|---|
| C | 41.14 | 42.56 | 42.48 |
| H | 3.95 | 4.51 | 4.54 |
| N | 17.13 | 17.32 | 17.34 |
| Cl | 26.02 | 19.46 | 26.02 |

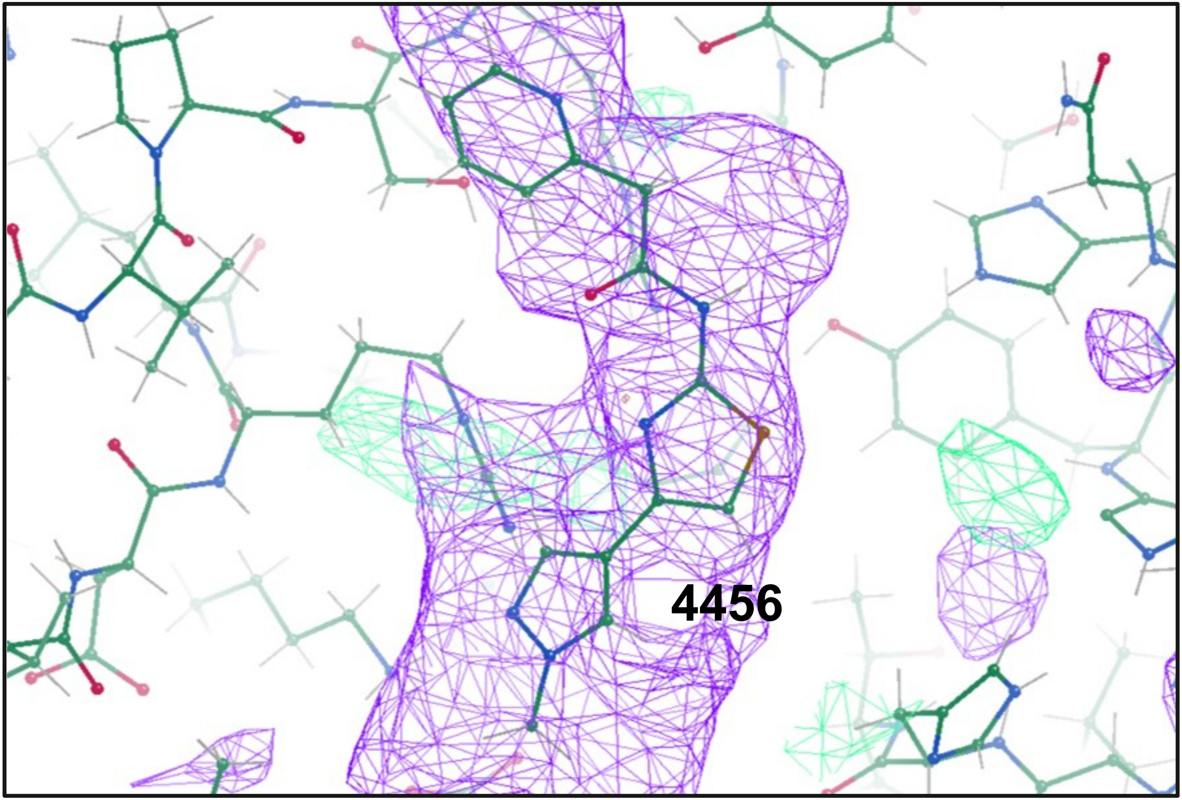
4456

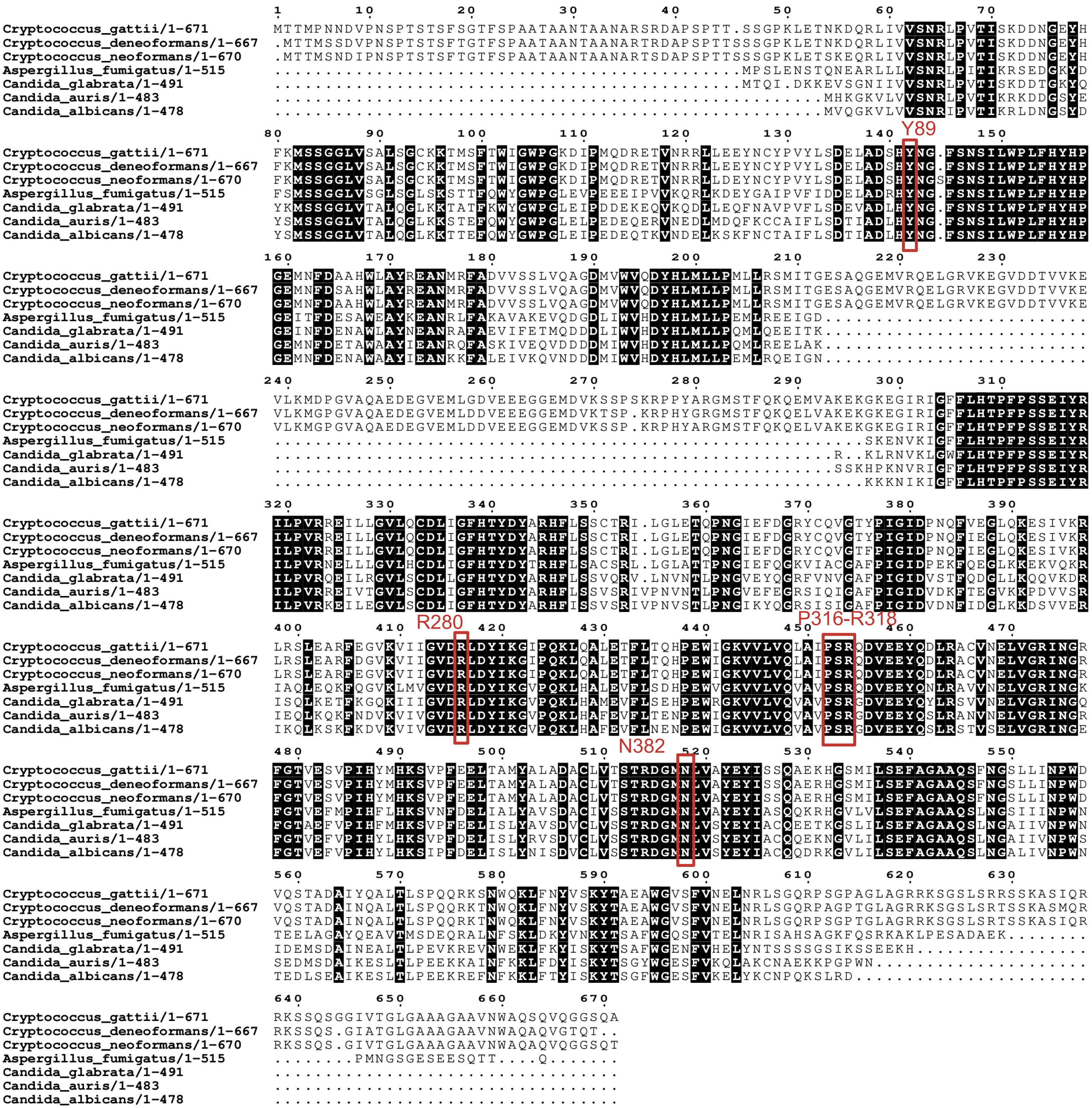

Cryptococcus_gattii/1-671
Cryptococcus_deneoformans/1-667
Cryptococcus_neoformans/1-670
Aspergillus_fumigatus/1-515
Candida_glabrata/1-491
Candida_auris/1-483
Candida_albicans/1-478
Y89
R280
P316-R318
N382

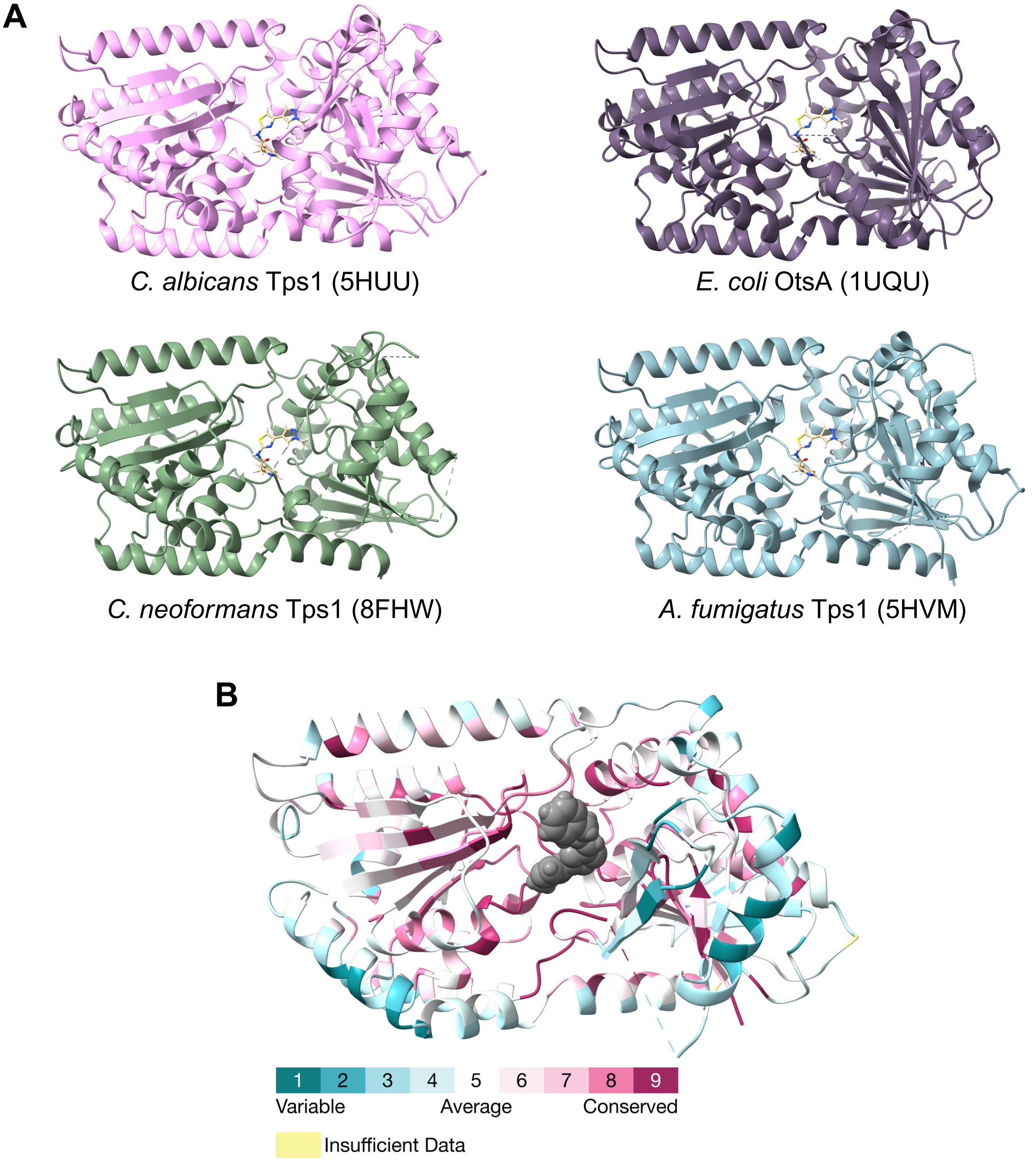
A
C. albicans Tps1 (5HUU)
E. coli OtsA (1UQU)
C. neoformans Tps1 (8FHW)
A. fumigatus Tps1 (5HVM)
B
1
2
3
4
5
6
7
8
9
Variable
Average
Conserved
Insufficient Data

A

| Fungal Species | $MIC_{80}$ (mM) |
|---|---|
| *C. albicans* | ND |
| *C. auris* | > 13 |
| *C. glabrata* | ND |
| *C. neoformans* | ND |
| *C. deneoformans* | > 13 |
| *C. gattii* | ND |

B

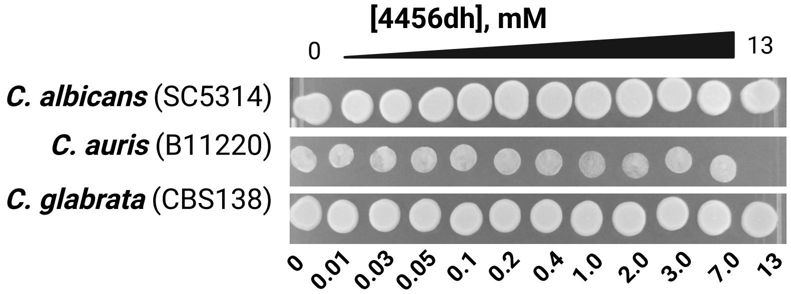


C

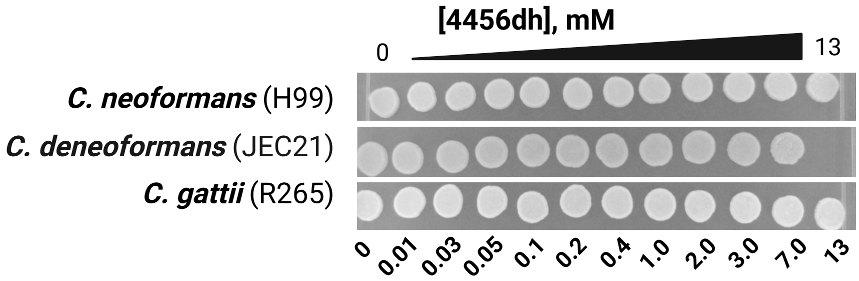

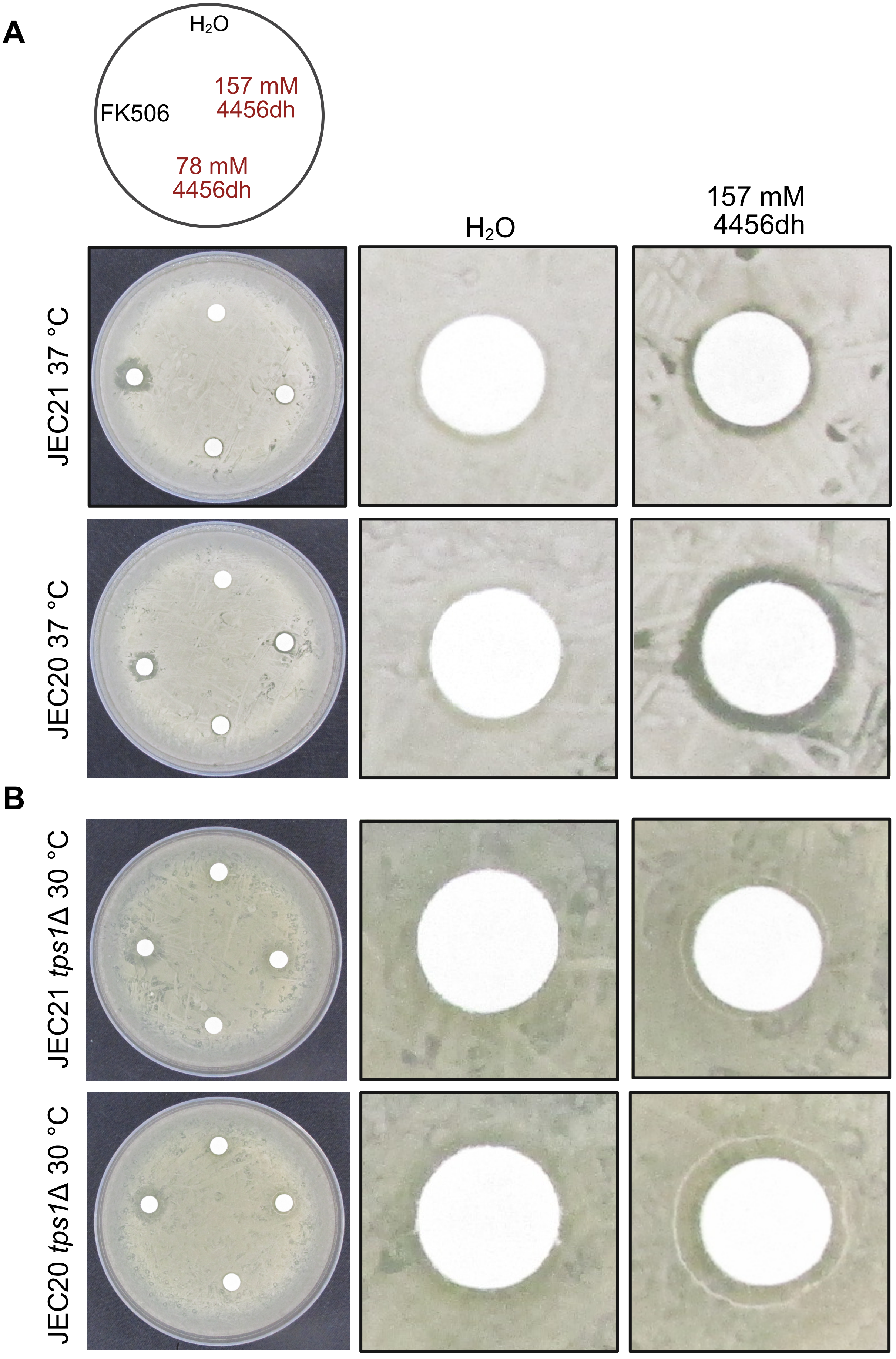
A
$H_2O$
157 mM
4456dh
FK506
78 mM
4456dh
$H_2O$
157 mM
4456dh
JEC21 37 °C
JEC20 37 °C
B
JEC21 *tps1*Δ 30 °C
JEC20 *tps1*Δ 30 °C

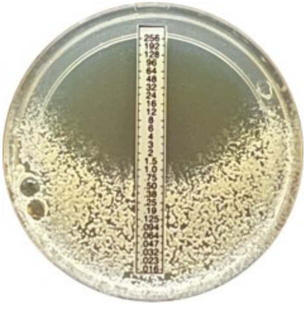
No 4456dh

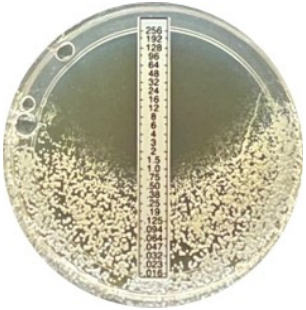
0.08 mM 4456dh

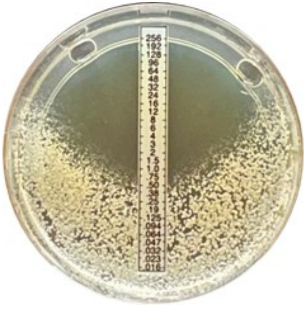
0.16 mM 4456dh

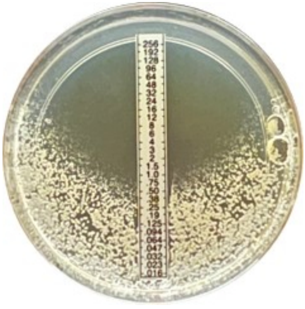
0.65 mM 4456dh

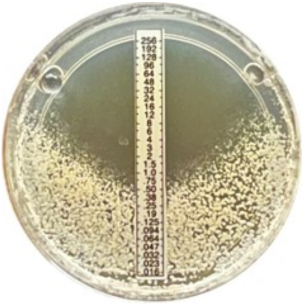
0.31 mM 4456dh

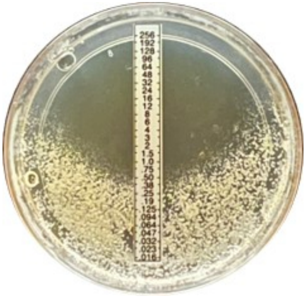
1.25 mM 4456dh

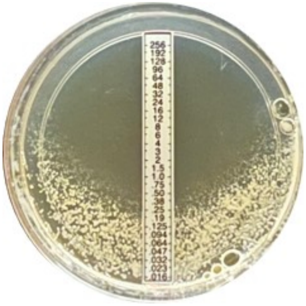
2.5 mM 4456dh

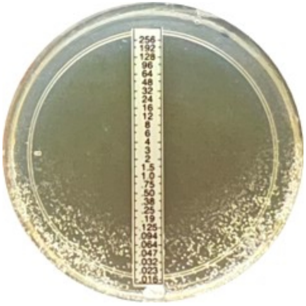
5 mM 4456dh

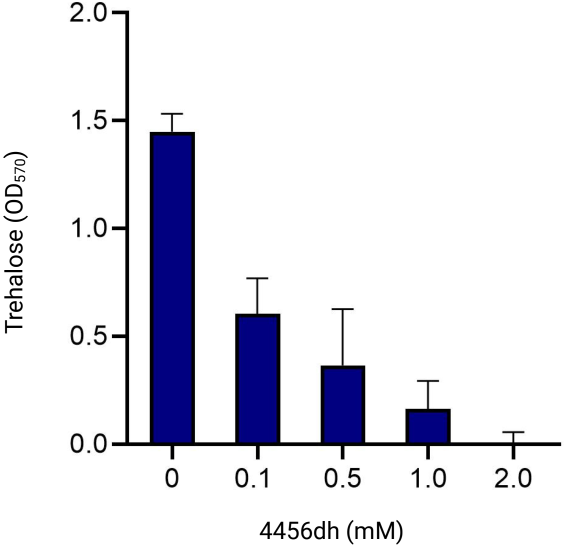

Trehalose ($OD_{570}$)
2.0
1.5
1.0
0.5
0.0
0
0.1
0.5
1.0
2.0
4456dh (mM)

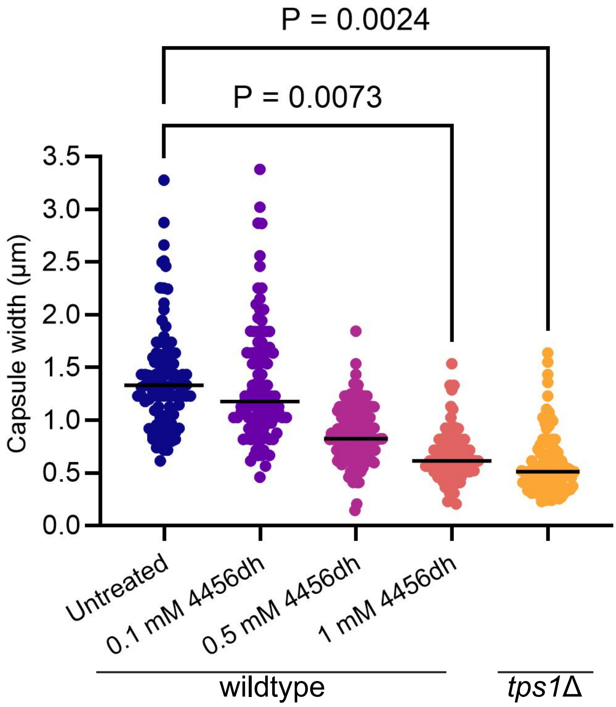

P = 0.0024
P = 0.0073
Capsule width (µm)
3.5
3.0
2.5
2.0
1.5
1.0
0.5
0.0
Untreated
0.1 mM 4456dh
0.5 mM 4456dh
1 mM 4456dh
wildtype
tps1Δ

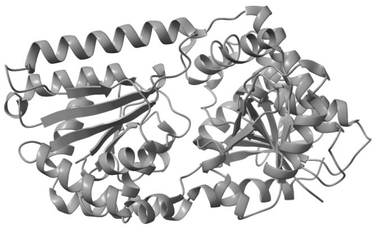

*C. albicans* Tps1 (5HUU)

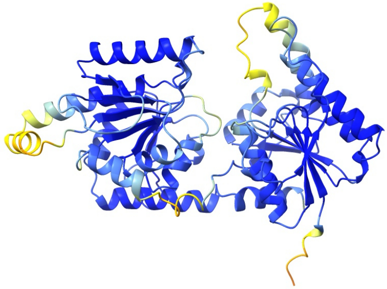

Q59LF2 Alpha-1,3/1,6-mannosyltransferase Alg2

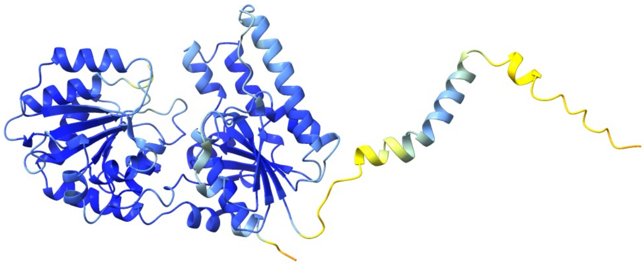

Q59Q79 Chitobiosyldiphosphodolichol beta mannosyltransferase

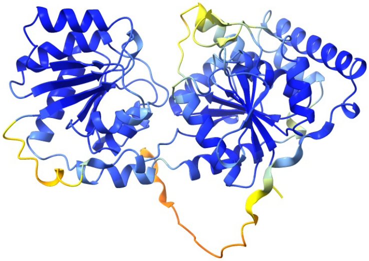

Q5A6R7 Glcnac-Pi Synthesis protein
